# Tunable biomimetic materials elaborated by ice templating and self-assembly of collagen for tubular tissue engineering

**DOI:** 10.1101/2023.08.30.555553

**Authors:** Isabelle Martinier, Florian Fage, Alshaba Kakar, Alessia Castagnino, Emeline Saindoy, Joni Frederick, Ilaria Onorati, Valérie Besnard, Abdul I. Barakat, Nicolas Dard, Emmanuel Martinod, Carole Planes, Léa Trichet, Francisco M. Fernandes

## Abstract

Synthetic tubular grafts currently used in clinical context fail frequently, and the expectations that biomimetic materials could tackle these limitations are high. However, developing tubular materials presenting structural, compositional and functional properties close to those of native tissues remains an unmet challenge. Here we describe a combination of ice templating and topotactic fibrillogenesis of type I collagen, the main component of tissues’ extracellular matrix, yielding highly concentrated yet porous tubular collagen materials with controlled hierarchical architecture at multiple length scales, the hallmark of native tissues’ organization. By modulating the thermal conductivity of the cylindrical molds, we tune the macroscopic porosity defined by ice. Coupling the aforementioned porosity patterns with two different fibrillogenesis routes results in a new family of materials whose textural features and the supramolecular arrangement of type I collagen are achieved. The resulting materials present hierarchical elastic properties and are successfully colonized by human endothelial cells and alveolar epithelial cells on the luminal side, and by human mesenchymal stem cells on the external side. The results reported here demonstrate the relevance of the proposed straightforward protocol, likely to be adapted for larger graft sizes, to address ever-growing clinical needs such as peripheral arterial disease or tracheal and bronchial reconstructions.

## 1. Introduction

In mammals, tubular tissues mediate macroscopic transport mechanisms in the vascular, respiratory, digestive and urogenital systems, ensuring that the contents of the lumen — gases, liquids or semi-solids — travel through the body without uncontrolled leakage to the surrounding environment. Because an important part of their function is to ensure the separation between different compartments in the body, tubular tissues cope with drastic biochemical gradients between the luminal and adventitial sides, the inside and outside of the tubes, respectively. In addition, variable pressure, strain at rest and contractile movements require specific mechanical behavior both under static and cyclic solicitation modes. Taken together, these constraints provide cues to understand the multiscale architecture that characterizes tubular tissues as well as the spatial distribution of the cells that populate them. Until now, the intrinsic complexity of these tissues has hindered the elaboration of biomimetic materials usable as tubular tissue grafts. The materials currently grafted in clinical practice to replace sections of tubular tissues meet the single purpose of providing mechanical support to the replaced tissue and fail in most of the other functions. Mimicking the native tissues’ complexity — in composition, structure and function — holds the potential to change drastically the materials available to engineer arteries, tracheas and other critical tubular tissues.

In general, cells displaying an epithelial phenotype line the luminal side of tubular tissues whereas different cell types of mesenchymal phenotype — fibroblasts, smooth muscle cells, or others, depending of the specific tissue in question — populate the wall of the tissue. In addition to the nature and spatial distribution of cells, tubular tissues display hierarchical arrangement of the extracellular matrix (ECM) components at several length scales, dictating their mechanical properties to a large extent. Type I collagen, the major component of the connective tissues in mammals, is widely responsible for the multiscale mechanical properties of native tissues. Locally, it provides the mechanical support upon which cells adhere, migrate and proliferate. At the macroscopic level, and together with cells and other components of the ECM, it provides the tissues with their viscoelastic response to the vast range of mechanical loading to which tubular tissues are subjected and ensures the dimensional stability of the tissues.

Moreover, *in vitro* as *in vivo*, type I collagen can be degraded and synthesized by different cells, providing the kind of dynamic environment that synthetic counterparts cannot provide. The preceding arguments confirm the relevance of type I collagen as the central component to develop scaffold-based grafts^[1]^ and to recapitulate native tissue-like architectures and fibrillar structures, thought to be key for attaining adequate mechanical properties and cell colonization^[2]^.

Although particularly complex to manipulate in non-denaturing conditions and at relevant physiological concentrations^[3]^, several techniques have been proposed to elaborate collagen-based tubular materials. Weinberg and Bell proposed a gel-based approach to develop vascular models by casting a suspension of cells in a collagen solution inside a tubular mold^[4]^. The resulting gel was further compacted though a cell-mediated process to reach materials that have been shown to be effective in preclinical studies, with functional restoration and vascular tissue recapitulation of the three different layers^[5]^. However, these materials required mechanical reinforcement by a synthetic sleeve for successful application. Cell-free strategies have equally been proposed to shorten fabrication times. Li *et al*. have reported a strategy based on drying and crosslinking collagen to modify the elastic properties of the tubular constructs^[6]^. In a different approach, dense collagen sheets obtained after plastic compression^[7]^ as well as composite matrices combining dried collagen gels and elastin^[8]^ were rolled around a cylindrical mandrel to obtain tubular materials. In the latter case the obtained vascular grafts demonstrated mechanical properties close to those of native tissues, with encouraging results obtained *in vivo*^[8]^. Despite these results, these techniques fail to produce materials that display both the fibrillar motifs of native tissues and the macroporosity required for successful cell colonization. Materials with accessible pores promote cell colonization^[9]^ inside the material and favor homeostasis due to an enhanced diffusion of nutrients and waste. Self-standing porous collagen materials can be obtained by lyophilization of collagen solutions followed by physical^[10]^ or chemical^[11]^ cross-linking. This approach was adapted by Koens *et al*. to provide tubular porous materials displaying three layers^[12]^, but one month after implantation thrombus formation was observed and associated to the porosity of the luminal wall. Collagen-based porous materials can also be obtained by means of biological textile approaches, which were pioneered by Cavallaro *et al*., with use of extruded, dehydrated and cross-linked collagen threads^[13]^. Electrochemically aligned collagen (ELAC) fibers were also used to knit tubular grafts. While the mechanical properties were satisfactory, the elevated pore size required an additional step of collagen electrospinning on the luminal surface while still requiring a cell-mediated production step of 6 to 12 weeks, followed by yarn fabrication^[14]^. These results highlight the importance of precise control over graft topography for successful graft implantation. Indeed, a *non-porous* luminal surface is required to foster the formation of epithelium or endothelium on the internal wall, while an oriented *porous* structure in the external part of the material would allow for effective cell colonization from the adjacent tissues.

Here, we introduce a new strategy based on ice templating to tackle the elaboration of materials that are fully composed of type I collagen, the main protein of the ECM, harness the principal mechanical and functional features of native tubular tissues and exhibit controlled macroporosity to favor cellularization. Ice templating (equally named freeze casting or ice-segregation-induced self-assembly), a technique initially developed for the elaboration of porous ceramics^[15]^ and adapted to produce scaffolds based on biopolymers^[16,17]^, offers the possibility to precisely tune the materials’ porosity by playing on different parameters such as solution composition, freezing kinetics, and temperature gradient^[18]^. Coupling ice templating to topotactic fibrillogenesis — a technique to induce the formation of native tissue-like collagen fibrils during thawing — enables self-assembly of type I collagen into biomimetic constructs while conserving the macroscopic patterns defined by the ice templating technique^[9,19]^. During freezing, the formation of ice crystals induces an increase in concentration of the collagen in solution, due to the insolubility of most solutes in ice^[20]^. The local concentration of collagen attained by this process reaches the same range of those observed in native tissues. The increase in concentration during freezing coupled to the lyotropic behavior of collagen in solution gives rise to highly organized domains, which, upon fibrillogenesis, recapitulate the multiscale hierarchical architecture of collagen in native tissues. Compared to other methods, this strategy enables obtaining materials with a controlled and hierarchical structure at different length scales by means of a straightforward protocol, likely to be adapted for larger scale graft production. In this work, we propose to use this technique to provide tubular materials applicable for tissue engineering purposes. In particular, we explored the effects of processing parameters such as the thermal conductivity of the cylindrical molds and fibrillogenesis routes on the textural and mechanical properties of the biomimetic tubular materials. In addition to the structural characterization we have performed *in vitro* colonization studies with a variety of cell lines to understand the effect of the materials’ processing routes on the colonization by human umbilical vein endothelial cells (HUVECs), adenocarcinomic human alveolar basal epithelial cells (A549), as well as human mesenchymal stem cells (hMSC).

The materials reported here combine, for the first time, the hierarchical organization found in native ECM — from the molecular scale up to the tissue level — using type I collagen, the mechanical performance in the range of native tissues as well as the capacity to be colonized by endothelial, epithelial and mesenchymal cell types. These findings open an exciting pathway towards the development of new biomimetic tubular tissue grafts that promise to find applications in pathologies such as peripheral arterial disease (PAD) or tracheal and bronchial reconstructions, for which the currently available materials fail.

## 2. Results

### 2.1. Controlled ice growth determines the macroscopic features of biomimetic tubular tissues

To elaborate collagen tubular materials *via* ice templating we have developed a series of tubular molds that were filled with type I collagen solution (40 mg.mL^-1^) and subsequently plunged in liquid nitrogen at a controlled velocity (Figures S1 & 1A). The difference between the thermal conductivity of the inner and outer parts of the mold enables the generation of a variety of thermal gradients across the solution and, as a consequence, the control of ice crystal growth. This simple approach allows the adjustment of both the size and the orientation of the ice crystals within the frozen monolith (Figure 1A). When exposed to a cryogenic liquid such as liquid nitrogen, insulating materials composing the mold delay the nucleation and growth of ice in the collagen solution, whereas thermally conductive materials promote faster nucleation and growth. The different materials used to study the effect of the thermal conductivity, λ, on the scaffold structure of the tubular matrices include aluminum (λ = 237 W.m^-1^.K^-1^), brass (λ = 110 W.m^-1^.K^-1^), stainless steel (λ = 45 W.m^-1^.K^-1^) and acrylonitrile butadiene styrene (ABS) (λ = 0.17 W.m^-1^.K^-1^). When the outer mold material displays lower thermal conductivity than the inner part of the mold, the freezing events proceed from the luminal surface towards the outer surface. At the initial moments of the freezing process, due to supercooling, a disordered zone appears close to the internal layer. When the solidification front reaches a steady state, ice crystals grow following the direction of the thermal gradient. Meanwhile, as the insulating outer mold is directly in contact with the cooling bath, ice also nucleates at its inner surface, but delayed in comparison to the luminal side, resulting in two frozen regions facing each other.

**Figure 1.**
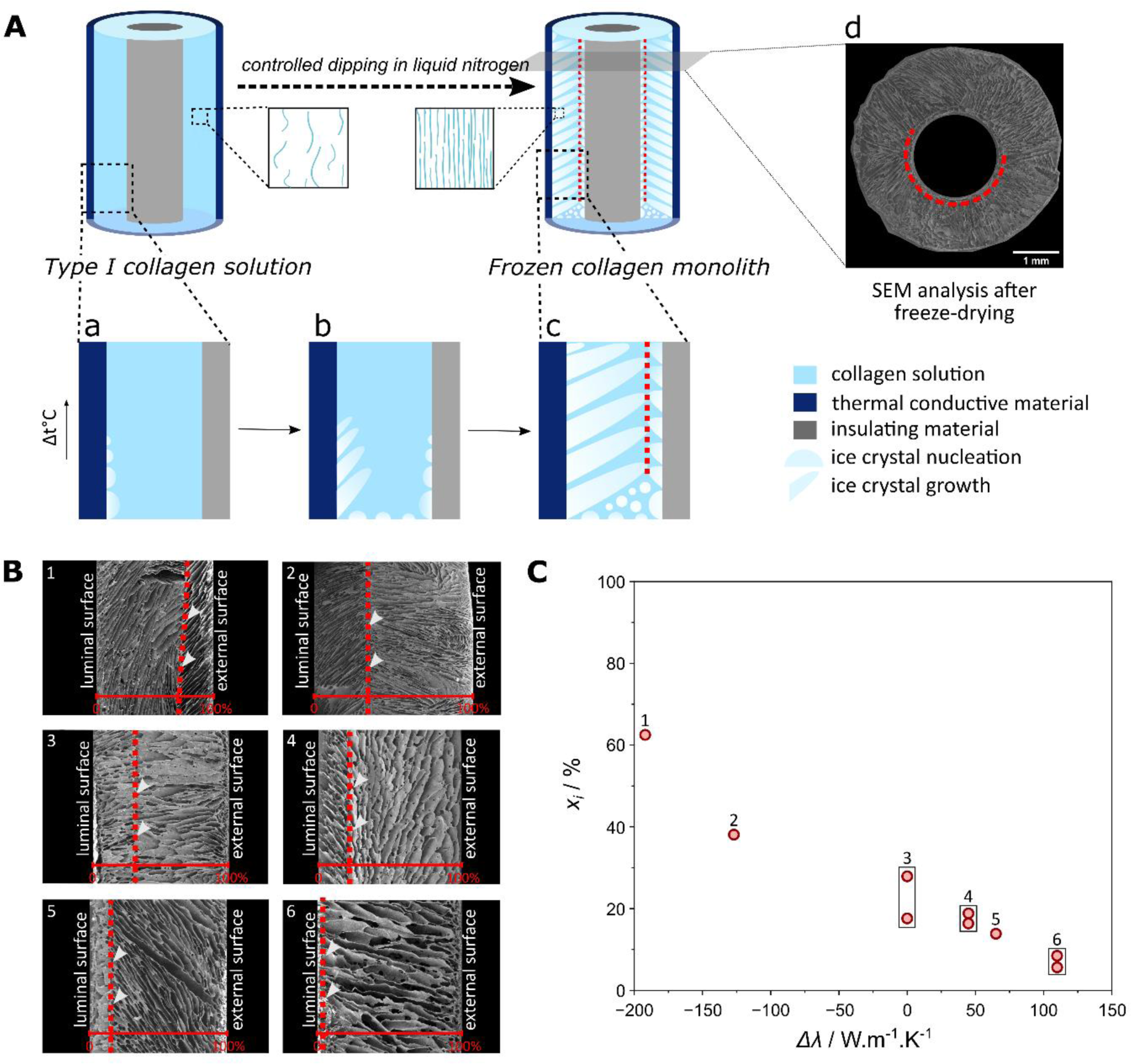
The internal texture of type I collagen materials obtained by ice templating is determined by the thermal conductivity difference between the inner and outer parts of the freezing molds. (**A**) Scheme depicting the ice nucleation and growth events in collagen-based tubular materials leading to the formation of an interface inside the walls of the tubes. SEM analysis of transverse sections of a selected material after freeze-drying showing the concentric position of the interface, highlighted with a red dotted line. (**B**) SEM images of longitudinal sections of freeze-dried collagen materials. The red dotted lines illustrate the position of the interface between the luminal (left) and external (right) surfaces. (**C**) Influence of the difference in molds’ thermal conductivities on the position of the interface formed in between the two ice growth regions. Numbers 1 to 6 refer to the SEM longitudinal sections shown in B for specific pairs of materials (displayed in Sup. Table 1). The interface position is defined as the percentage of the distance to the lumen with respect to the total wall thickness of the matrix.

Observed after lyophilization, the porous structure of the collagen matrix is the fingerprint of the ice crystals formed during the cooling process. On both sides, lamellar pores are formed and converge towards an interfacial zone highlighted with dotted lines on SEM transverse section images after lyophilization (Figure 1A). The diffqerent pairs of molds, as well as their thermal conductivity difference (Δλ = λ_*out*_ − λ_*in*_), are displayed in Sup. Table 1. SEM images of the longitudinal cross-sections of the collagen matrices after freeze-drying fabricated with a variety of mold materials are shown in Fig. 1B (top).

For the freezing conditions chosen here (dipping speed and mold geometry), and for the materials mentioned above, the position of the interface, *x*_*i*_, scales linearly with Δλ and can be adjusted between 5.6% and 62.5% of the tube wall as observed in Fig. 1B and Fig. 1C. The relevance of this interface is multiple since it is expected to play a role in the materials’ mechanical performances, the permeability towards cells and fluids as well as in the regulation of the cells’ migration path.

### 2.2. Controlling the dimensions and orientation of macropores in collagen scaffolds using ice

The ability to control the size and the orientation of pores in collagen scaffolds is expected to play an important part in cellular adhesion, proliferation and access to nutrients as well as to promote — or hinder — cell migration within the material. We have selected two different pairs of materials — for the inner and outer parts of the mold — that illustrate the wide range of porous structures attainable through ice templating of biomimetic tubular scaffolds. Materials fabricated with an outer conductive material (aluminum) and an inner insulating one (ABS) are referred to as ***Cond***, while scaffolds prepared with both parts of the mold composed of insulating material (ABS) are referred to as ***Ins***. These conditions yield Δλ = 236.83 *W*. *m*^−1^. *K*^−1^ and Δλ = 0 *W*. *m*^−1^. *K*^−1^, respectively.

Samples were ice templated under equivalent boundary conditions (dipping speed, air and bath temperature) using both mold pairs (*Cond* and *Ins*). After lyophilization, longitudinal and transversal sections of the scaffolds were imaged by SEM to observe the orientation and dimensions of the pores, as shown in Fig. 2.

**Figure 2.**
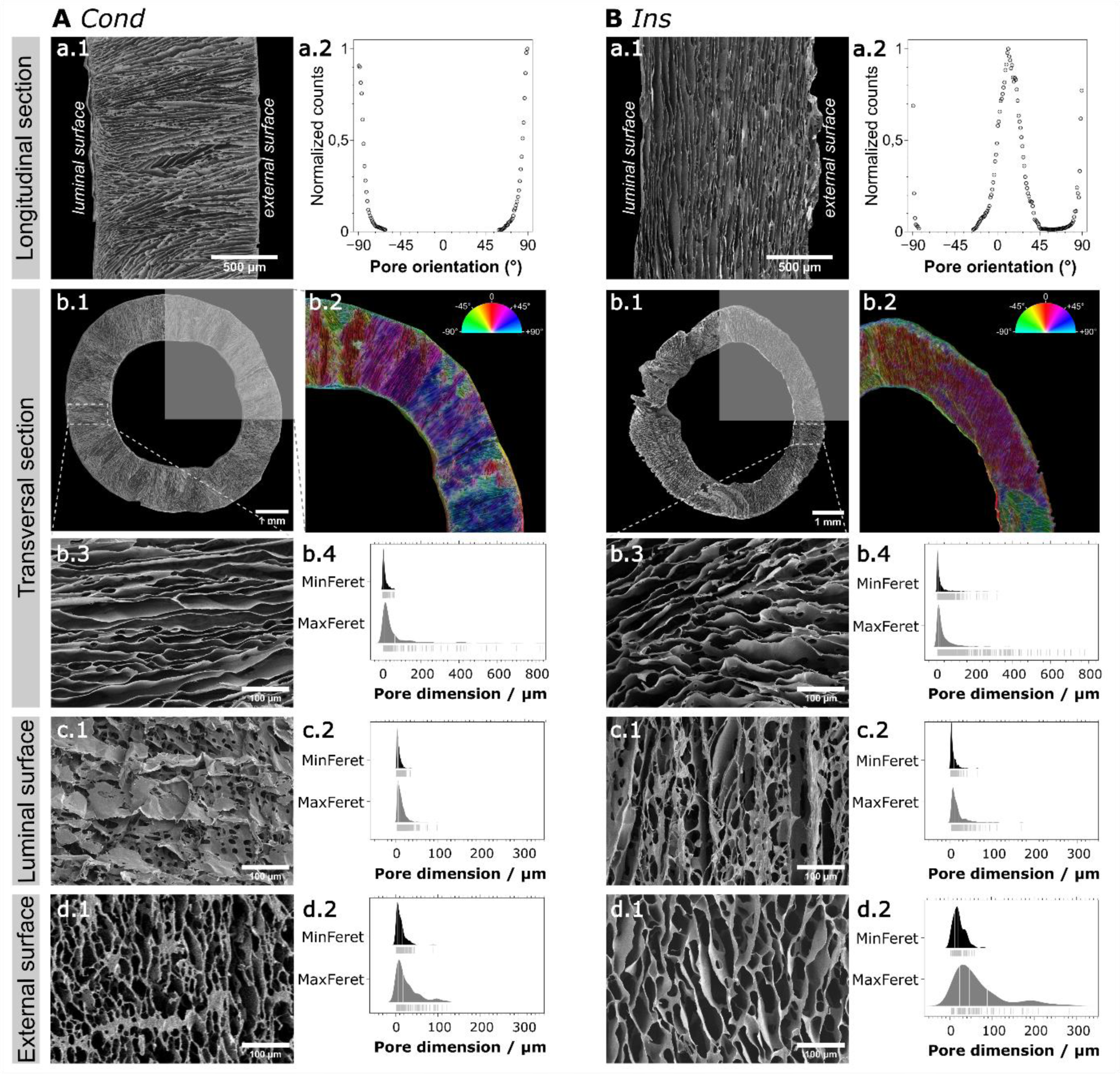
Textural properties of collagen scaffolds obtained with pairs of materials differing in their conductive properties. (**A**) Scaffolds fabricated using an outer conductive mold and an inner insulating mold (*Cond*). (**B**) Scaffold fabricated with insulating materials for both outer and inner molds (*Ins*). a.1: longitudinal cross-section of A and B, imaged by SEM. a.2: analysis of pore orientation on longitudinal cross-section, with respect to the main longitudinal axis of the scaffold. b.1: transversal cross-section of the scaffolds, imaged by SEM. b2: analysis of pore orientation on transversal cross-section, with indication of angle orientation based on color code mapping. b.3: high magnification of the zone defined with dotted lines on b.2 images. b.4: analysis of Minimum Feret (MinFeret) and Feret diameters (MaxFeret) of the pores on transversal sections of A and B. c1: luminal (inner) surface of the tubular scaffolds, imaged by SEM. c.2: analysis of MinFeret and MaxFeret diameters of the pores on the luminal surface of each scaffold. d.1: external (outer) surface of the tubular scaffolds, imaged by SEM. d.2: analysis of MinFeret and MaxFeret diameters of the pores on the external surface of A and B.

Pore orientation analysis on SEM longitudinal sections (Figure 2A & B.a.1), showed that the pores are aligned perpendicular (*c.a*. 90°) to the longitudinal axis of the tube for the ***Cond*** freezing conditions (Figure 2A.a.2). Conversely, ***Ins*** conditions favor pores oriented at 12° with respect to the longitudinal axis of the tube (Figure 2B.a.2). These results are in accordance with the fact that the outer conductive material favors a radial thermal gradient within the collagen solution. At the initial moment of freezing, ice nucleates in contact with the outer mold, and grows towards the lumen, as confirmed by the SEM observation of the transversal cross-section of ***Cond*** (Figure 2A.b.1). ***Ins*** freezing conditions favor the axial thermal gradient imposed by the dipping process, yielding a different pore organization, as shown on both SEM longitudinal and transversal cross-sections (Figure 2B.b.1).

Pore orientation was analyzed on a quarter of each tubular material cross-section (highlighted in grey on Figs. 2A & B.b.1) using the OrientationJ plugin in FIJI^[21]^ (Figure 2A & B.b.2). In ***Cond*** materials, the continuous change of orientation of the pores, observed by the various colors, corresponds to a radial organization. On the contrary, ***Ins*** materials feature pores that are organized in coherent domains with a major orientation consistent with the longitudinal axis of the tube. Higher magnification images on other specific parts of the tubular walls confirm this observation (Figure 2A & B.b.3). The striking difference in the internal porosity of these materials is determined solely by the thermal gradient imposed by the mold walls.

The size of the pores observed transversally and at the luminal surface appears to be independent from the molds’ thermal conductivity difference. In fact, pores exhibit comparable dimensions with a median Feret diameter of 25.75 µm and 9.77 µm in ***Cond*** scaffolds, and 20.62 µm and 11.36 µm in ***Ins*** scaffolds, for transversal sections and luminal surfaces respectively (Table 1). However, pore formation and dimensions at the external surfaces significantly relate to the thermal conductivity of the outer mold. The median Feret diameter and percentage of surface accessible porosity are both lower in ***Cond***, with a Feret value almost 3 times higher for ***Ins*** (45.83 µm vs 17.32 µm), and a percentage of surface accessible porosity 1.5 times higher for ***Ins*** scaffolds than for ***Cond*** scaffolds (48% vs 63%) (Fig. 2A & B.d,12 and Table 1). These differences may be explained by the ice nucleation conditions imposed by the various pairs of mold materials. In particular, since freezing occurs later under ***Ins*** conditions, it is likely that a wider supercooling zone is formed before freezing, whereas this zone should be minimal in an earlier freezing process (*Cond*). All the preceding textural features were observed in dry conditions, after ice templating and lyophilization. The key challenge is to maintain the level of control over the porosity obtained by the ice templating process after fibrillogenesis (*i.e*. after inducing the self-assembly of collagen into a hydrated fibrillar hydrogel). This would ensure that the biochemical cues from native fibrillar collagen could be combined with the physical environment created by the freezing conditions.

**Table 1.**
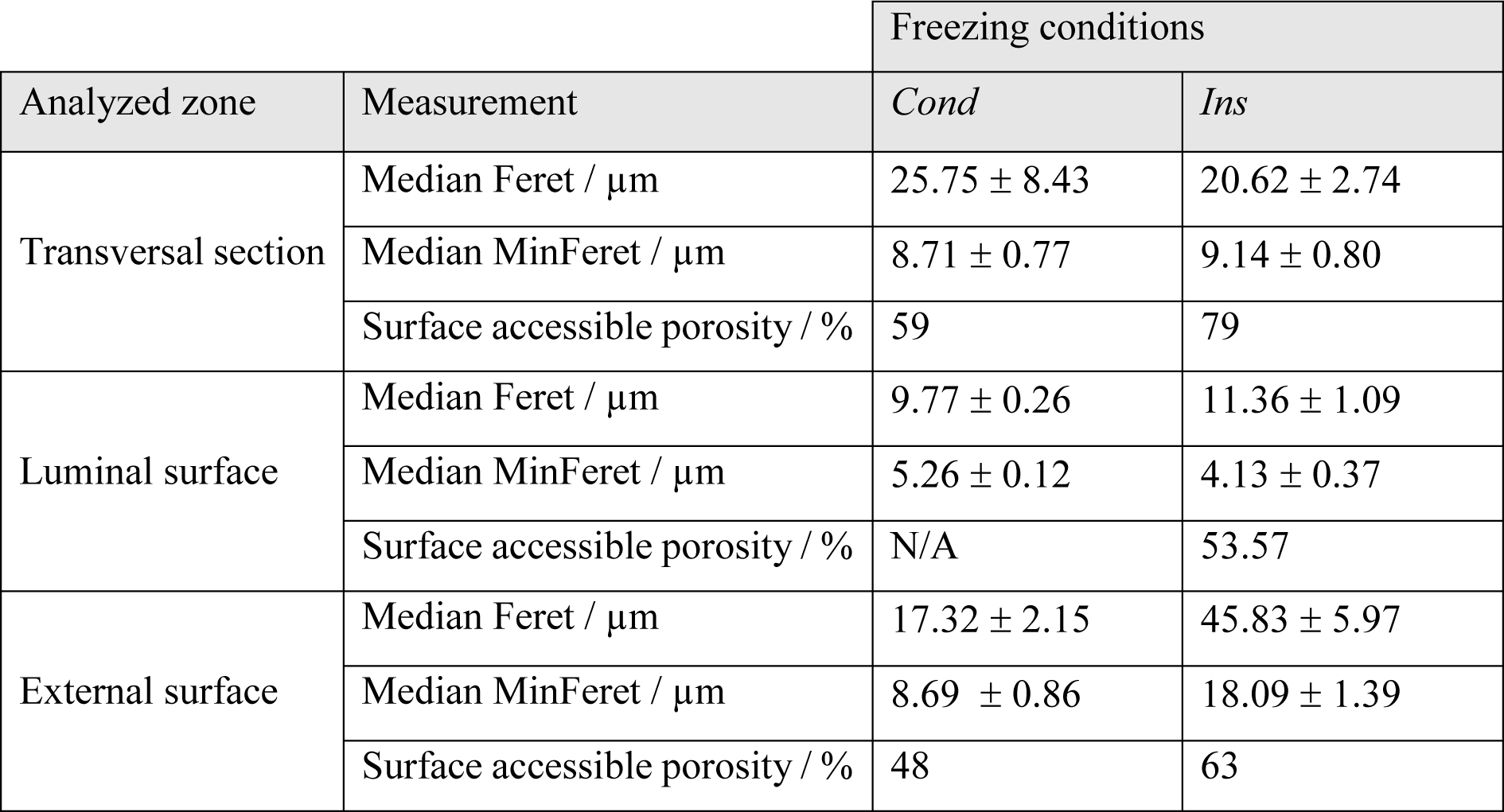
Dimensions of the scaffold pores for various sections and surfaces, after ice-templating and freeze-drying.

### 2.3. Influence of the topotactic fibrillogenesis pathway on the porous structure

Topotactic fibrillogenesis refers to the ability to induce fibrillogenesis by a pH rise in a collagen-based material during thawing while simultaneously keeping the macroscopic textural characteristics developed during freezing^[19]^. To maintain the shape and dimensions of the tubular scaffolds as well as their macroporous texture after ice-templating, two fibrillogenesis processes were conducted here, one corresponding to the gas-phase ammonia process previously described^[9,19]^ (Figure 3A), while the other uses a concentrated phosphate buffer saline (PBS (10X)) solution (Figure 3B). In both cases, the challenge is to use a single step process to remove the ice crystals formed during ice templating while inducing fibrillogenesis — resulting in an insoluble type I collagen gel. When in direct contact, ammonia vapors decrease the freezing point of ice. By exposing the frozen sample to ammonia vapors at 0 °C we ensured that the only ice crystals able to melt were those in direct contact with NH_3_ vapors. Therefore, near the ice crystals being thawed, collagen molecules were exposed to a pH value above its isoelectric point, inducing their self-assembly into native tissue-like fibrils. Changing the fibrillogenesis route towards a liquid-solid interface rather than a vapor-solid interface is challenging. To ensure that the competition between the kinetics of the ice thawing and fibrillogenesis were sufficiently balanced to obtain a self-supported fibrillar material, we have performed the process at a lower temperature (-3°C).

**Figure 3.**
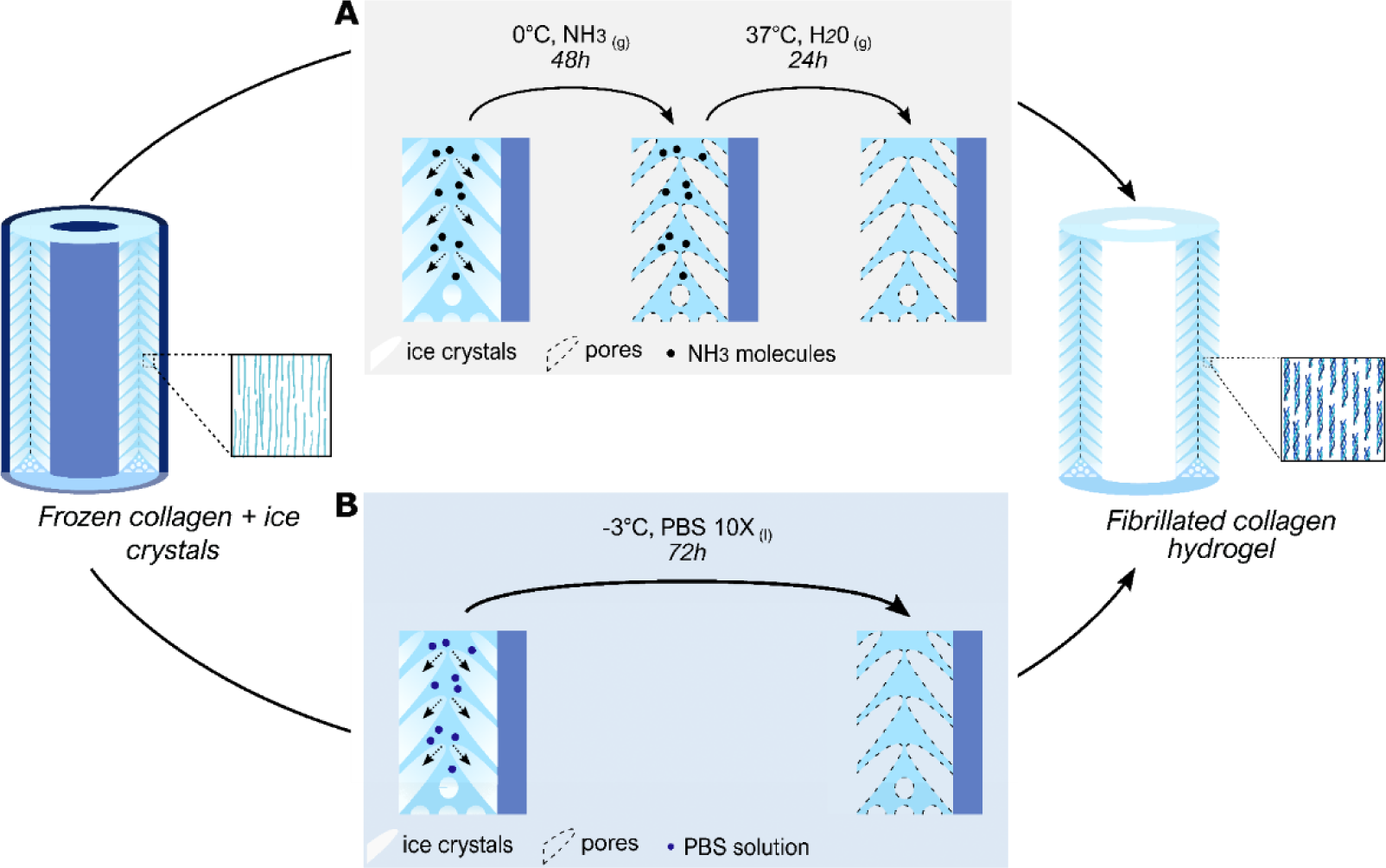
Fibrillogenesis routes applied after ice-templating of the collagen scaffolds and removal of the outer mold. (**A**) Vapor-solid phase process using ammonia vapors at 0°C for 48h, followed by desorption in a water-vapor bath for 24h. (**B**) Liquid-solid based process using PBS (10X) in solution at -3°C for 72h. Both procedures are followed by a 2-week PBS (5X) bath at room temperature to secure the fibrillogenesis. The resulting self-standing hydrogels are highly concentrated in fibrillated type I collagen with tunable porosity.

Confocal images of the scaffolds confirm the ability to induce the topotactic fibrillogenesis of collagen scaffolds *via* the ammonia gas-phase route (noted ***Cond-*** or ***Ins-*NH_3_**). Pores formed during freezing remain open and interconnected, and the orientation imposed during the ice-templating process is maintained (Figure 4A.1.a & 2.a). Conversely, the liquid-phase fibrillogenesis pathway induces either the collapse (Figure 4B.1.a) or a strong reduction of the pore size (Figure 4B.2.a) for ***Cond-*PBS** and ***Ins-*PBS** samples, respectively.

**Figure 4.**
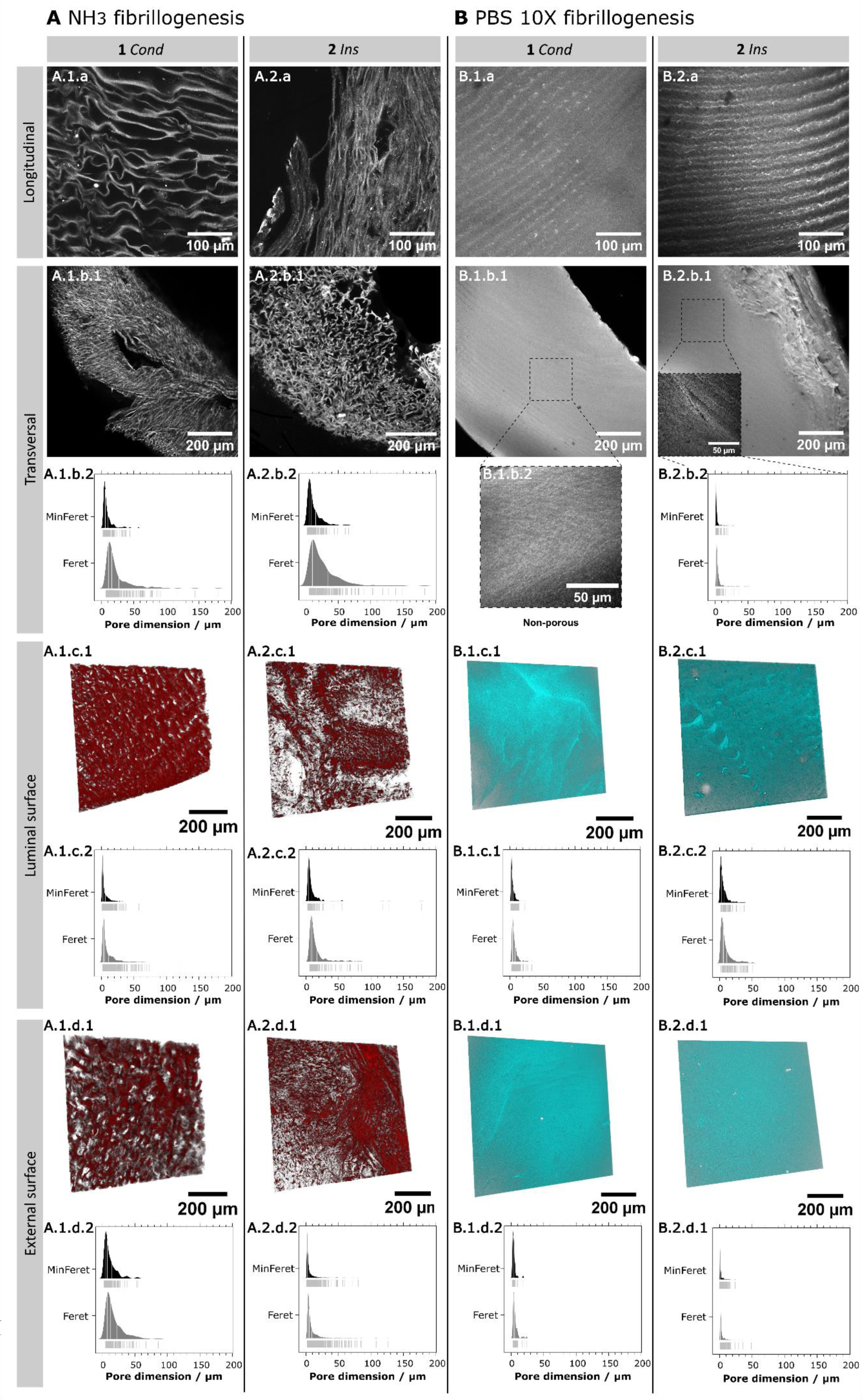
Textural properties of collagen scaffolds after topotactic fibrillogenesis, by gas-phase or liquid-phase pathways, imaged by confocal microscopy. (**A**) Scaffolds fibrillated by the gas-phase method, based on ammonia vapors at 0°C. (**B**) Scaffolds fibrillated by the liquid-phase method, based on a buffer solution of PBS (10X) at -3°C. A.1. A.2. B.1. B.2: (a) longitudinal cross-sections of the scaffolds (b.1) transversal cross-sections of the scaffolds and (b.2) the analysis of Min and MaxFeret diameters of the transversal pores for each. (c.1) 3D-rendering of the luminal (inner) layer (100-200 µm) of the tubular scaffolds and (c.2) the analysis of Min and MaxFeret diameters of the pores for each. (d.1): 3D-rendering of the external (outer) layer (100-200 µm) of the tubular scaffolds and (c.2) the analysis of Min and MaxFeret diameters of the pores for each.

***Cond-*NH_3_** and ***Ins-*NH_3_** feature transversal pore sizes (median Feret diameter (Feret) x median minimum Feret diameter (MinFeret)) of (16.12 ± 1.17 µm × 6.31 ± 0.39 µm) and (18.76 ± 1.57 µm × 8.05 ± 0.64 µm), respectively (Figure 4 b.1-2 for A.1 and A.2, and Table 2). Compared to ice-templated samples that were lyophilized, the pore size decreases during fibrillogenesis with a reduction of the Feret diameter of 37% and 9% for ***Cond*** and ***Ins*** conditions, respectively.

**Table 2.** Dimensions of the scaffold pores for various sections and surfaces, after ice-templating and topotactic fibrillogenesis. When available, values are given with associated standard deviation.

| Analyzed zone | Measurement | Freezing and fibrillogenesis conditions |  |  |  |
| --- | --- | --- | --- | --- | --- |
|  |  | NH<sub>3</sub> |  | PBS (10X) |  |
|  |  | Cond | Ins | Cond | Ins |
| Transversal section | Median Feret / µm | 16.12 ± 1.17 | 18.76 ± 1.57 | N/A | 3.78 ± 0.27 |
|  | Median MinFeret / µm | 6.31 ± 0.39 | 8.05 ± 0.64 | N/A | 1.78 ± 0.14 |
|  | Median aspect ratio | 3.04 ± 0.10 | 2.33 ± 0.08 | N/A | 1.84 ± 0.05 |
|  | Surface accessible porosity / % | 24 | 34 | N/A | 31 |
| Luminal surface | Median Feret / µm | 10.41 ± 2.49 | 16.00 ± 2.25 | 5.18 ± 0.26 | 5.98 ± 0.60 |
|  | Median MinFeret / µm | 5.56 ± 1.51 | 7.82 ± 1.18 | 3.16 ± 0.16 | 3.32 ± 0.37 |
|  | Median aspect ratio | 2.03 ± 0.07 | 1.97 ± 0.97 | 1.91 ± 0.06 | 1.47 ± 0.04 |
|  | Surface accessible porosity / % | 53 | 21 | <1 | 5 |
| External surface | Median Feret / µm | 5.04 ± 0.27 | 19.35 ± 3.28 | 4.78 ± 0.54 | 1.43 ± 0.12 |
|  | Median MinFeret / µm | 2.68 ± 0.15 | 10.22 ± 1.54 | 3.43 ± 0.35 | 0.79 ± 0.07 |
|  | Median aspect ratio | 1.99 ± 0.02 | 1.78 ± 0.10 | 1.33 ± 0.09 | 1.46 ± 0.02 |
|  | Surface accessible porosity / % | 50 | 28 | <1 | 2 |

As ammonia reaches the surface of ice crystals, a competitive process takes place between the thawing of ice crystals induced by cryoscopic depression of the ammonia-ice interface and the fibrillogenesis of collagen. The swelling of the collagen walls is a direct consequence of ice thawing being favored over collagen fibrillogenesis. In such a case, a partial dilution of the collagen walls in the liquid water generated by the ice melting occurs — before the pH change induces the formation of the collagen gel — leading to smaller pore size in the hydrogel than in the dried scaffold. We hypothesize that the lower pore diameter reduction for ***Ins*-NH_3_** is due to its unidirectional pore structure, which may favor faster diffusion of ammonia molecules and subsequent fibrillogenesis of collagen. On the other hand, the isotropic texture of ***Cond*** samples may induce slower diffusion of ammonia and lead to faster ice melting and subsequent increased dilution of collagen towards the pore space. This observation is equally confirmed by the proportion of pores giving access to the inner wall in Figure A.1.c.1 and A.2.c.1, where the respective pore coverage for ***Ins-*NH_3_** and ***Cond-*NH_3_** decreases from 35% to 24%.

In contrast, PBS (10X)-fibrillated scaffolds demonstrate reduced pore accessibility in transversal plane (Figure 4B.1.b.1 and B.2.b.1), and regardless of the ∼20 µm periodic artifacts due to the sample cutting process the method fails to retain pore orientation (Figure 4B.1.a and B.2.a). ***Cond*-PBS** samples are devoid of pores at the observable length scale. At higher magnification, a few pores can be observed at the surface of the transversal sections (Figure 4B.1.b.2) closest to the lumen in ***Ins-*PBS**. In the center, small pores can be witnessed in between the formed fibrils (Figure 4B.2.b.1), with a median Feret diameter of 3.78 ± 0.27 µm and an accessible pore surface of 30.5% (Figure 4B.2.b.2). The conditions imposed by the fibrillogenesis in PBS (10X) favor the melting of ice crystals prior to the self-assembly of collagen. The architecture of ice crystals plays an important role in the preservation of their shape and the subsequent porosity. Unidirectional pore structures such as those found in ***Ins*** favor faster diffusion of the PBS ions, slightly shifting the kinetics in favor of collagen self-assembly, associated with pore preservation.

The pore size on the luminal surface is not altered by the gas-phase fibrillogenesis process (Figure 4 c.1-2, d.1-2 for A.1 and A.2). Pores in the lumen present Feret diameters of 10.41 µm and 16.0 µm for respectively ***Cond-*NH_3_** and ***Ins-*NH_3_** (Table 2), versus 9.77 µm and 11.36 µm in lyophilized scaffolds (Table 1). Pores of the external surface are 2 to 4 times smaller than those of the lyophilized scaffolds, with a mean Feret diameter of 5.04 µm and 19.35 µm for respectively ***Cond*** and ***Ins***, compared to 17.32 µm and 45.83 µm in dry state. However, the degree of surface accessible pores differs from what was observed for lyophilized scaffolds: ***Cond*** provides the highest pore coverage with 53% at the lumen and 50% at the external surface, compared to 21% and 28% for ***Ins***. We can hypothesize that the mismatch between the conservation of pore size on one hand and the reduction of surface accessible pores on the other hand may be attributed to a contraction of all pores following fibrillogenesis. The smallest pores observed after freeze-drying may vanish or become too small to be detected, while the largest pores may be reduced due to a slight collagen wall swelling between the dry and hydrogel forms. In contrast, fibrillogenesis in PBS (10X) leads to a significant reduction of the pore size at the luminal and external surfaces (Figure 4 c.1-2, d.1-2 for B.1 and B.2). At the lumen, pore size is reduced by a factor of ∼2 with a Feret diameter shifting from 9.77 µm to 5.18 µm in ***Cond*-PBS** and from 11.36 µm to 5.98 µm in ***Ins*-PBS**. On the external surface, Feret pore diameter is respectively 4x and 32x smaller in ***Cond*-PBS** and ***Ins*-PBS**, shifting from 17.32 µm to 4.78 µm, and from 45.83 µm to 1.43 µm. More importantly, the degree of accessible pores decreases most notably in ***Cond*-PBS** where the proportion of pores granting access to the inner wall on the lumen and external surfaces amounts to less than 1% of the total surfaces. In ***Ins*-PBS**, 5% and 2% of the luminal and external surfaces are covered by pores, corresponding to a 90% size reduction in comparison to lyophilized scaffolds. PBS (10X) route results in the formation of more rounded pores, characterized by a mean aspect ratio ranging between 1.58 and 1.8, in contrast to NH_3_ fibrillogenesis that yields values ranging from 2.05 to 2.30. As in any temperature-dependent process, the conditions of topotactic fibrillogenesis determine the final textural characteristics of the scaffolds. The gas-phase route favors the self-assembly of the collagen molecules that stiffen the walls surrounding the ice crystals and subsequently melt the ice crystals to reveal the pores. The resulting materials correspond thus to a fibrillar gel sample whose texture is the fingerprint of the ice crystals formed during ice templating. Conversely, the liquid-phase fibrillogenesis favors first the thawing of ice-crystals, resulting in a dilution of the collagen molecules initially concentrated in the interstices and a partial or complete loss of the texture defined by ice. In the latter case, shifting the kinetics in favor of self-assembly can enhance the preservation of the porous network. Anisotropic structures — generated by using inner and outer molds of equivalent thermal conductivity as in ***Ins*** — lead to faster diffusion rate of the fibrillation media associated with increased likelihood of partially keeping the pores and their interconnectivity. However, the evolution of some textural features — such as the reduction of pore size — induced by liquid route topotactic fibrillogenesis can present advantages. Specifically, the pore dimensions on the luminal surface of tubular constructs obtained through PBS (10X) topotactic fibrillogenesis closely resemble those observed in the tracheal basal membrane, which display an average diameter of 1.76 µm^[22]^. In this sense, the liquid-phase fibrillogenesis route described here can be seen as a supplementary tool for fine-tuning the porosity of the materials and matching the textural characteristics of a given native tissue.

### 2.4. Influence of the fibrillogenesis pathway on the collagen molecular and supramolecular arrangement

During freezing, collagen molecules in solution are forced to concentrate in the interstitial space created in between ice crystals. Because collagen in solution behaves as a lyotrope, molecular crowding of collagen can lead to the formation of biological analogues of liquid crystals^[23]^. In specific cases, collagen crowding can also lead to local prefibrillar states, as suggested by Gobeaux *et al*.^[24]^. Although the results cited above were obtained at room temperature and there are no reports of such mesophases or prefibrillar arrangements near water freezing conditions, it is likely that these semi-ordered states exist in solution close to 0°C. To probe the molecular arrangement of collagen molecules in the obtained fibrillar materials, we examined 200 µm-thick transversal sections of the scaffolds under an optical microscope equipped with cross-polarizers. Each of the samples display birefringent signal along the axis defined by the bisections of the quadrants formed by the analyzer and polarizer positioned at 0° and 90°, respectively (Figure 5 1-2.a for A and B). The ***Cond-*NH_3_** sample features alternating patches of birefringent and extinction zones throughout the entire thickness of the wall (Figure 5A.1.a). These birefringent domains, in the tens of µm across, are dispersed homogeneously throughout the whole thickness of the tubular wall. The ***Ins-*NH_3_** sample exhibits a remarkably different behavior, with the presence of a large birefringent domain at the scale of the full thickness of the tube wall. The difference between the birefringence signals among these scaffolds highlights the importance of the freezing conditions in obtaining diverse textures, even when employing the same fibrillogenesis protocols. Samples fibrillated in PBS (10X) also display a collective birefringent signal spanning the entire thickness of the samples, suggesting the existence of contiguous organized domains of collagen molecules throughout the sample walls. In general, the birefringent signal found at the luminal and external walls suggests a greater degree of order in these zones, possibly resulting from the concentration of collagen molecules in the initial moments of ice templating.

**Figure 5.**
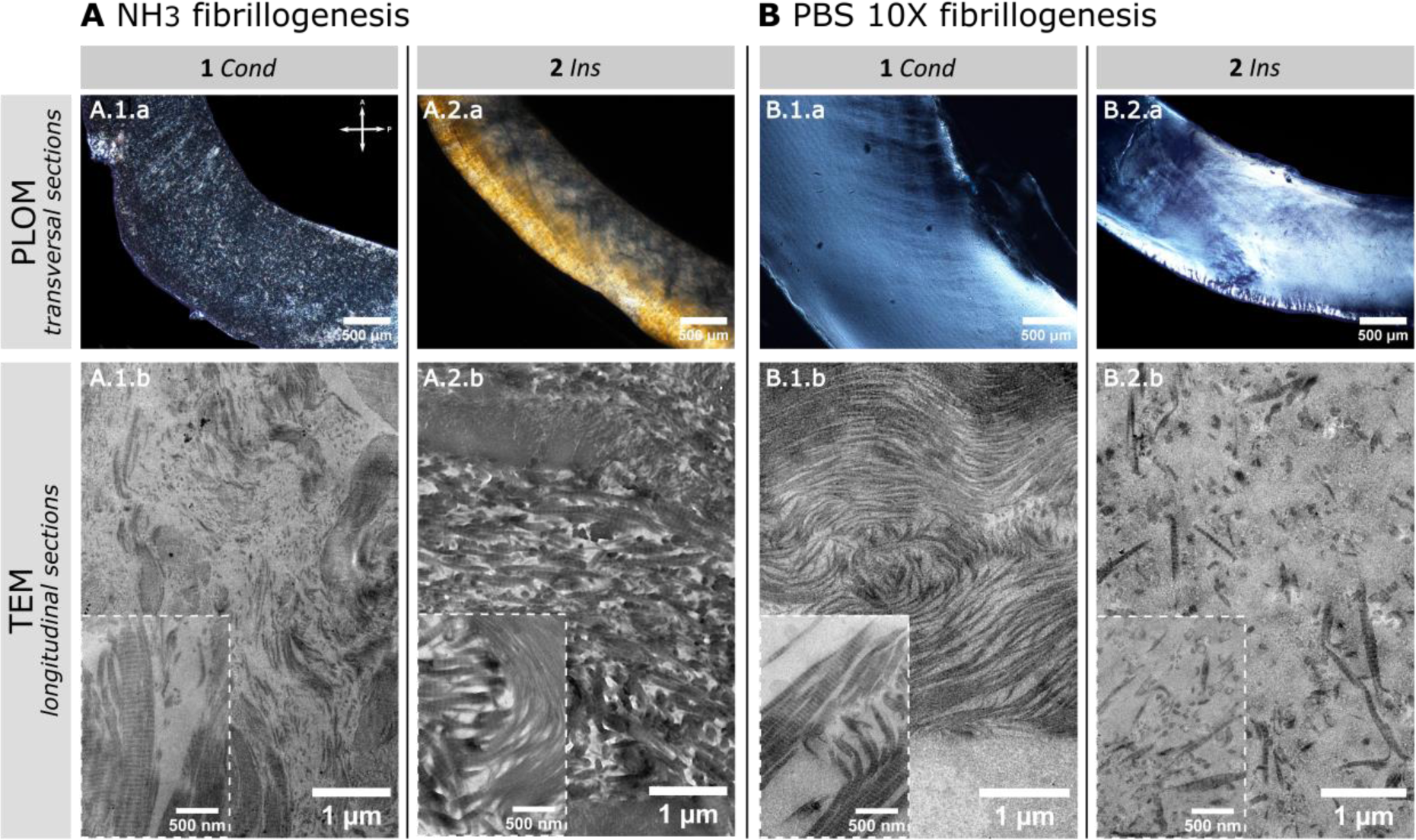
Influence of the processing conditions on the supramolecular organization of collagen in the scaffolds. Materials were fibrillated using ammonia (**A**) or PBS (10X) (**B**), with either conductive (**1**) or insulating (**2**) outer molds. (A & B.1.a, A & B.2.a): under cross-polarizers oriented at 0°, the transversal sections all exhibit a birefringent signal, characteristic of liquid crystal-like ordered structures. Transmission electron micrographs of the materials (A & B.1.b, A & B.2.b). Insets display the typical D-banding striation found in the fibrils of each sample.

Ordered structures revealed by the birefringence signal emerge from two factors that are difficult to disentangle. On the one hand, molecular crowding can lead to the formation of mesophases, and in some case to the appearance of pre-fibrillar structures, able to produce birefringence signal^[24]^. On the other hand, since samples are exposed to physicochemical conditions meant to induce fibrillogenesis — ammonia vapors and PBS (10X) solution — even collagen solutions displaying an isotropic molecular arrangement may lead to fibrillar structures able to produce birefringent signal. At the observed scale here, polarized optical microscopy provides clues to understand the global orientation of the collagen molecules within the tubular samples, but is insufficient to describe the formation of biomimetic fibrillar domains.

Ultra-thin longitudinal sections of the scaffolds observed under TEM demonstrate the existence of fibrillar structures for each ice templating/topotactic fibrillogenesis condition, similar to the motifs found in the ECM of arterial tissue. Observed at higher magnification, fibrils present a D-banding pattern along their axis, with distances below the typical 67 nm striations due to their off-plane orientation with respect to the cross-section plane (Figure S3). Regardless of the observed striation distances, the existence of these features confirms the fact that both fibrillogenesis routes induce the formation of biomimetic fibrils. Exposed to ammonia vapors, collagen molecules form a dense network of small fibrils (Figure 5-A.1.b and A.2.b), both transversal and parallel to the observation plane. Fibrils in ***Cond-*NH_3_** are smaller and lie in a looser arrangement than those in ***Ins-*NH_3_**, revealing that a simple change of the thermal conductivity of the molds during ice-templating can affect collagen fibril formation. Fibrils in ***Ins-*NH_3_** form compact arched patterns, surrounded by denser regions. Conversely, PBS (10X) promotes the formation of a higher number of large striated fibrils, in less tightly packed domains. These results confirm the observations of the texture observed under the confocal microscope where the swelling during fibrillogenesis led to narrower — or absent — pores. We hypothesize that during the partial melting of the ice crystals, concentrated collagen solution in the interstitial space is rediluted, which increases the mobility of the molecules and enables the formation of a greater number of supramolecular structures. The high compaction of the fibril network in ***Cond-*NH_3_***, **Ins-*****NH_3_** and ***Cond-*PBS**, is similar to what is seen in native tissues, suggesting a local concentration that ranges between 50 and 200 mg.mL^-1[25]^. The diversity of fibril sizes and arrangement observed here could provide a valuable tool to elaborate materials capable of mimicking specific features of the ECM from a variety of tissues.

Although the preceding results confirm the relevance of our elaboration strategy in obtaining biomimetic materials that display structural and textural features of the ECM, they do not provide direct proof of the stabilization effect brought by the fibrillogenesis processes in cold conditions. We have measured the thermal stability of the different materials using Differential Scanning Calorimetry (DSC) to characterize the thermal denaturation processes of each sample type (Figure S4). Samples fabricated in **-PBS** are more stable, with a mean denaturation temperature (T_d_) 2.6°C higher than with **-NH_3_** (Table 3). Such a difference can be explained by the prevalence of larger and more mature fibrils in these samples, leading to higher stability. The structural variations induced by the conductive or insulating outer mold do not significantly alter the T_d_. In all four conditions, the denaturation temperature is considerably above that of non-fibrillar materials around 37°C.

**Table 3.** Thermal stability measurements of the materials demonstrate that PBS (10X) fibrillogenesis leads to a more stable collagen network by the increased denaturation temperature. No significant difference is noted between the materials fabricated with a conductive (***Cond***) or insulating (***Ins***) outer mold. Denaturation temperatures (T_d_) of the materials were measured by differential scanning calorimetry (DSC) (Figure S4).

|  | NH<sub>3</sub> fibrillogenesis |  | PBS (10X) fibrillogenesis |  |
| --- | --- | --- | --- | --- |
|  | *Cond* | *Ins* | *Cond* | *Ins* |
| $T_d \pm SD / ^\circ\text{C}$ | $48.73 \pm 3.41$ | $46.76 \pm 0.52$ | $49.55 \pm 1.09$ | $51.11 \pm 0.99$ |

Two major differences standout regarding the influence of the fibrillogenesis pathways on the collagen texture and microstructure. Ammonia treatment enables the maintenance of the porosity and of the collagen walls’ concentration. It leads to the formation of small fibrils, tightly packed in a complex 3D arrangement according to the TEM and PLOM images. PBS (10X) favors the thawing of ice, resulting in less concentrated collagen domains that display larger and more mature fibrils which increase the thermal stability. Regardless of the structure, it is evident that a modification of the physico-chemical parameters induces variations in the size and arrangement of fibrils. We offer here conditions that facilitate the stabilization of the liquid crystal-like mesophases, yielding materials that mimic the ECM of native tissues. These features are expected to promote *in vivo* cellularization following implantation given their similarities to the native cellular microenvironment.

### 2.5. Multiscale elastic properties of biomimetic tubular materials

The mechanics of biological tissues and those of their analogues are defined by — at least — two different length scales that correspond to the cell microenvironment and to the full tissue. We have characterized the elastic properties of the biomimetic tubular materials using AFM and quasi-static macroscopic traction tests to describe the material behavior under mechanical loading.

The resistance to macroscopic deformation in both the longitudinal and circumferential directions under quasi-static loading was evaluated for the four different types of obtained tubular materials (Figure 6A and 6B). In the longitudinal direction, ***Ins***-**NH_3_** samples exhibited the highest stiffness, with a Young’s modulus (E) of approximately 58 ± 39 kPa (mean ± SD). Conversely, the circumferential stiffness of ***Ins***-**NH_3_** samples, as well as the longitudinal and circumferential stiffnesses of ***Cond***-**NH_3_** samples, were notably lower, ranging between 6 and 10 kPa. These findings indicate that the matrices display anisotropic elastic properties under macroscopic loading, with greater rigidity observed along the longitudinal direction of ***Ins***-**NH_3_** samples compared to others. Under ***Ins*** conditions, the process of ice templating involves the uniaxial growth of ice crystals, which alignment is preserved through ammonia fibrillogenesis. The longitudinal orientation of pores explains the greatest Young’s modulus obtained for ***Ins***-**NH_3_**, and is coherent with superior elastic properties in the direction of crystal growth but poorer elastic properties in the transverse directions. In contrast, the thermal gradient imposed in ***Cond*** leads to radially oriented collagen walls and pores. As a consequence, the circumferential and longitudinal directions are perpendicular to the porous structure and show a relatively low Young’s modulus. The analogy to the elastic properties of composite materials, here with a collagen matrix populated by zero Young’s moduli inclusions — the pores — allows us to rationalize this behavior. One may invoke a Rule of Mixtures (ROM) behavior in case the pores and the walls are aligned with the traction direction, whereas an Inverse Rule of Mixtures (IROM)^[26]^ may explain the lower moduli when the traction is applied perpendicular to the main pore and wall directions.

**Figure 6.**
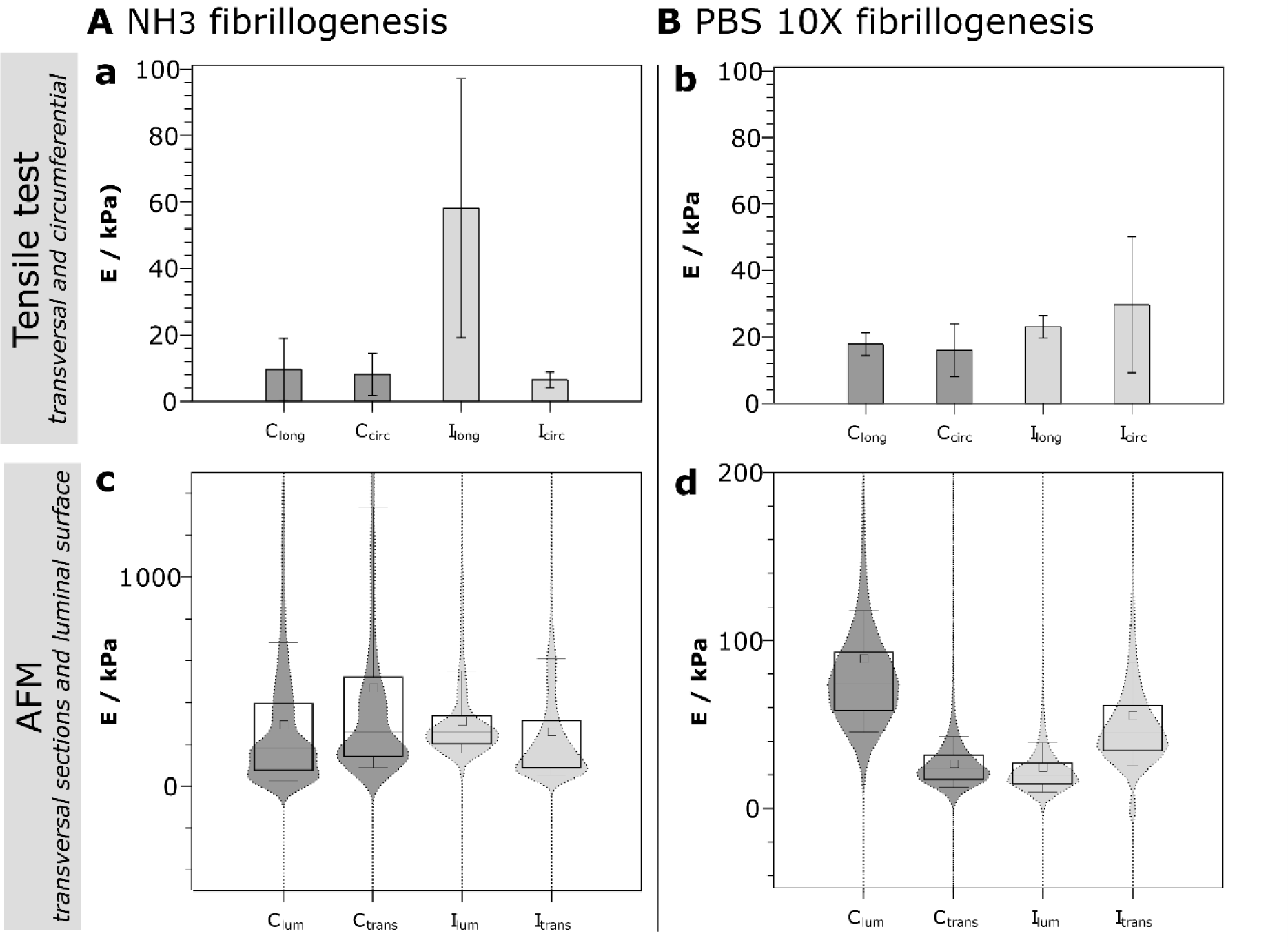
Influence of the processing conditions on the mechanical properties of the scaffolds. Materials were fibrillated using ammonia (**A**) or PBS (10X) (**B**), with either conductive (*Cond* - **C**) or insulating (*Ins* - **I**) outer molds. (a, b) Longitudinal (long) and circumferential (circ) traction Young’s moduli for each sample. Error bars represent standard deviations. (c, d) Luminal (lumen) and transversal (trans) compression Young’s moduli for each sample. Box plots represent the 25th to 75th percentile range of the modulus distribution, with the whiskers indicating the 10th to the 90th percentile range of the modulus distribution. The squares denote the mean values, and the straight lines represent the medians.

Contrary to the results obtained using NH_3_ as a fibrillogenesis medium, PBS (10X) (Figure 6B) provides the tubular materials with isotropic elastic properties. Regardless of the orientation of the pores initially developed through ice templating, elastic moduli remain close to 20 kPa. This behavior is ascribed to the fact that PBS (10X) favors faster ice melting over the self-assembly of collagen, resulting in poorly preserved porosity, small pore size and low anisotropy compared to NH_3_-fibrillated samples (Figure 4). Instead of having a composite material with highly distinct mechanical properties in the longitudinal and circumferential directions, as seen in NH_3_-fibrillated samples, we obtain a material with similar mechanical properties in both deformation directions. This difference in elastic properties depending on the direction of stress between the two types of fibrillogenesis could allow mimicking a large range of tissues with orthotropic properties (such as bone, cartilage, tendon…) as well as tissues with isotropic properties (adipose tissue, connective tissue, spongy bone…).

The elastic properties earlier discussed describe the behavior of the materials at the tissue level. However, their local elastic properties — at the cell length scale — are equally important to ascertain their relevance as tissue grafts. The local Young’s modulus was measured in compression on the luminal and transverse surfaces of -**PBS** and **-NH_3_** samples using an Atomic Force Microscope (AFM). Unlike the tensile measurements that provided the macroscopic stiffness of the samples, these local stiffness measurements focused on the stiffness of the walls within the samples. Results depicted in Figure 6 demonstrate that the **-NH_3_** samples exhibited a significantly higher average stiffness of approximately 300 kPa compared to the **-PBS** samples, which had an average stiffness of around 50 kPa. Furthermore, the ammonia-fibrillated samples exhibited a broader distribution, characterized by increased variability in the 25th and 75th percentile values when compared to the **-PBS** fibrillated samples. Additionally, the violin plots suggest the co-existence of two populations of elastic local environments for the **-NH_3_** samples, while the **-PBS** sample displays a Gaussian distribution indicative of a single population. Taken together, these results confirm the observations arising from the microscopy techniques (TEM and confocal microscopy) that suggested a partial dilution of the collagen walls during (10X) PBS-induced topotactic fibrillogenesis, whereas NH_3_ tends to preserve highly concentrated collagen domains in the internal walls of the scaffolds. The ability to tune the collagen concentration in our scaffolds through different fibrillogenesis protocols presents an interesting aspect. Cells are significantly impacted by the collagen concentration in their microenvironment^[27]^, directly influencing the stiffness and mechanical properties of the surrounding extracellular matrix, as demonstrated by our AFM experiments. This crucially affects cellular adhesion, migration, differentiation, and tissue development. Consequently, tailoring local collagen concentration with our different fibrillogenesis protocols could result in adapted substrates for grafting at different locations with diverse surrounding cell types.

### 2.6. In vitro cellularization of biomimetic tubular materials Endothelialization and epithelialization of the luminal surface of tubular scaffolds

Functional epithelia and endothelia play a key role in tubular tissues because they provide the semi-permeable character that mediates transport from the luminal to the adventitial sides of the tissues. Human alveolar basal epithelial cells (A549) and human umbilical vein endothelial cells (HUVECs) were seeded on the luminal side of the different materials to ascertain the effect of the topography and the fibrillogenesis pathway on their suitability to host cells. The cellularization was studied in time and compared to a simplified model of basal lamina (*i.e*. PDMS substrates coated with type IV collagen) as a control. Observations under confocal microscopy of immunostained cells demonstrate that luminal surfaces of all materials promote cellular proliferation after a few days of culture (Figure 7).

**Figure 7.**
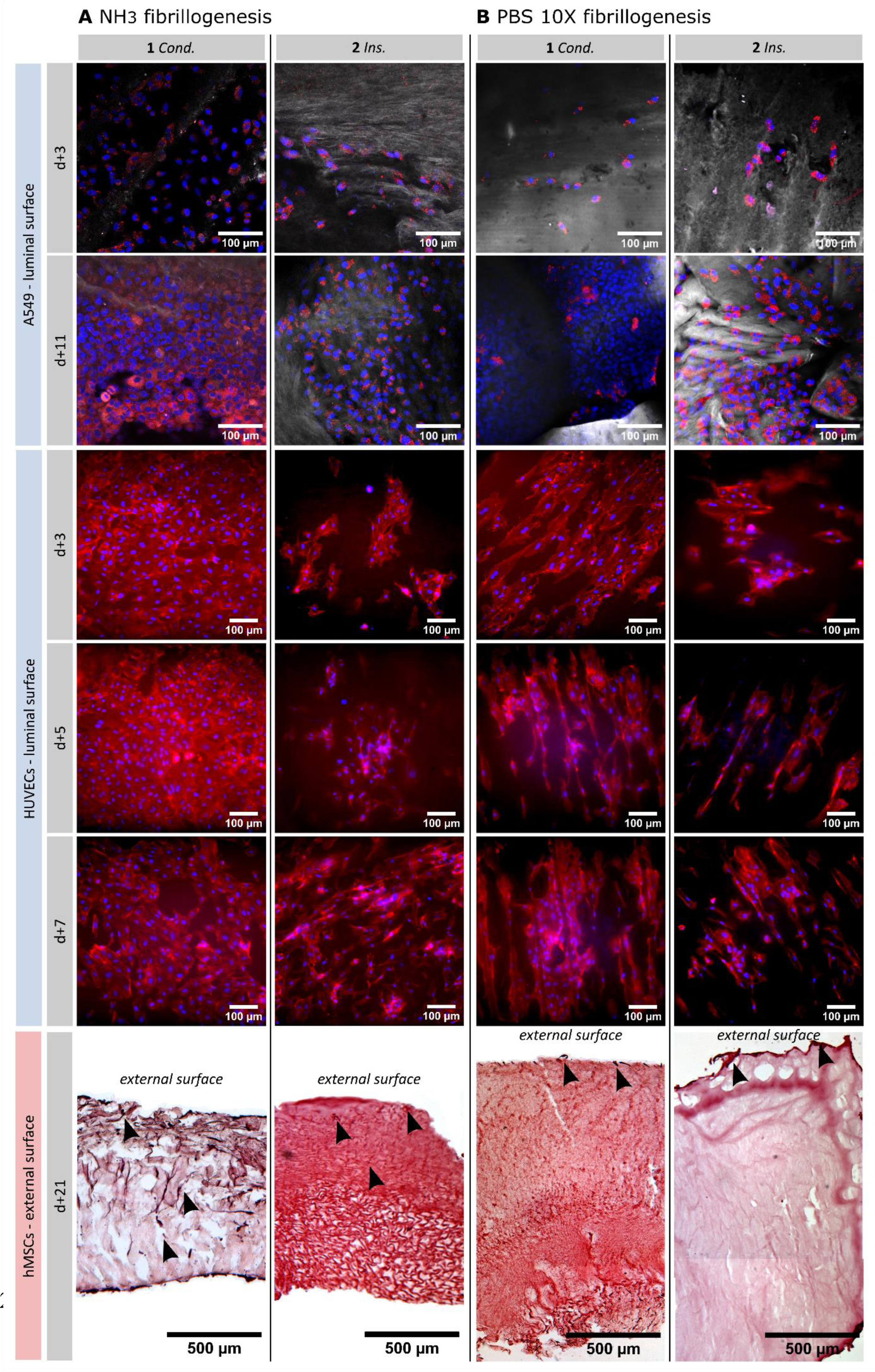
Cell colonization of biomimetic type I collagen materials for the studied mold materials and fibrillogenesis routes. Epithelium formation was studied with luminal seeding of two different cell lines, A549 cells (days 3 and 11) and HUVECs (days 3, 5 and 7). The ability of cells to colonize the adventitial part was demonstrated with hMSC and observed at day 21. Observations of epithelial cells were performed under confocal microscopy, with immunostaining of the nuclei (blue: DAPI), actin (red: phalloidin) and collagen (grey: FITC). Observations of the colonization by hMSC were performed under microscope on cryosections stained with hematoxylin (nucleus) and eosin (cytosol) dyes.

A549 display a homogenous covering of ***Cond*-NH_3_** surface at day 3 while the distribution of cells seems more erratic on other surfaces at that time, suggesting that ***Cond*-NH_3_** is more compatible for A549 adhesion. However, at day 11 for a given material, cell densities reach comparable levels in between the two fibrillogenesis and cell spatial distribution becomes more homogeneous, although ***Cond*** surfaces seem more suitable than ***Ins*** ones.

The trends of epithelial cellularization observed for these materials using A549 cells were equally confirmed by seeding HUVECs. As depicted in Fig. 7B, ***Cond*** surfaces generally provide a more favorable substrate for HUVECs adhesion than ***Ins*** surfaces with the cells exhibiting a more homogeneous distribution and more complete surface coverage regardless of the cell culture time and the fibrillogenesis pathway (with the exception of ***Cond-*PBS** at day 3). ***Cond-*NH_3_** cell density follows that of the biomimetic control at days 3, 5 and 7. At day 7 cell densities are steady compared to the previous time point, suggesting that confluence has been attained.

Materials prepared by PBS (10X) fibrillogenesis display slower cellularization kinetics as compared to materials fibrillated in ammonia. Despite the initial differences, the cell density in PBS (10X) fibrillogenesis materials does evolve positively over time as can be observed from Figs. 7 and 8. These results demonstrate that it is possible to direct the cell-material interactions by an adequate choice of the mold thermal conductivity parameters and the fibrillogenesis route. The highest density was obtained for ***Cond***, which coincides with the low porosity of the lumen previously observed under confocal microscopy. Combined with NH_3_ fibrillogenesis, ***Cond-*NH_3_** surfaces present the highest cell densities at early cell culture time points, but as mentioned earlier cell density appears to decrease once confluence is attained.

**Figure 8.**
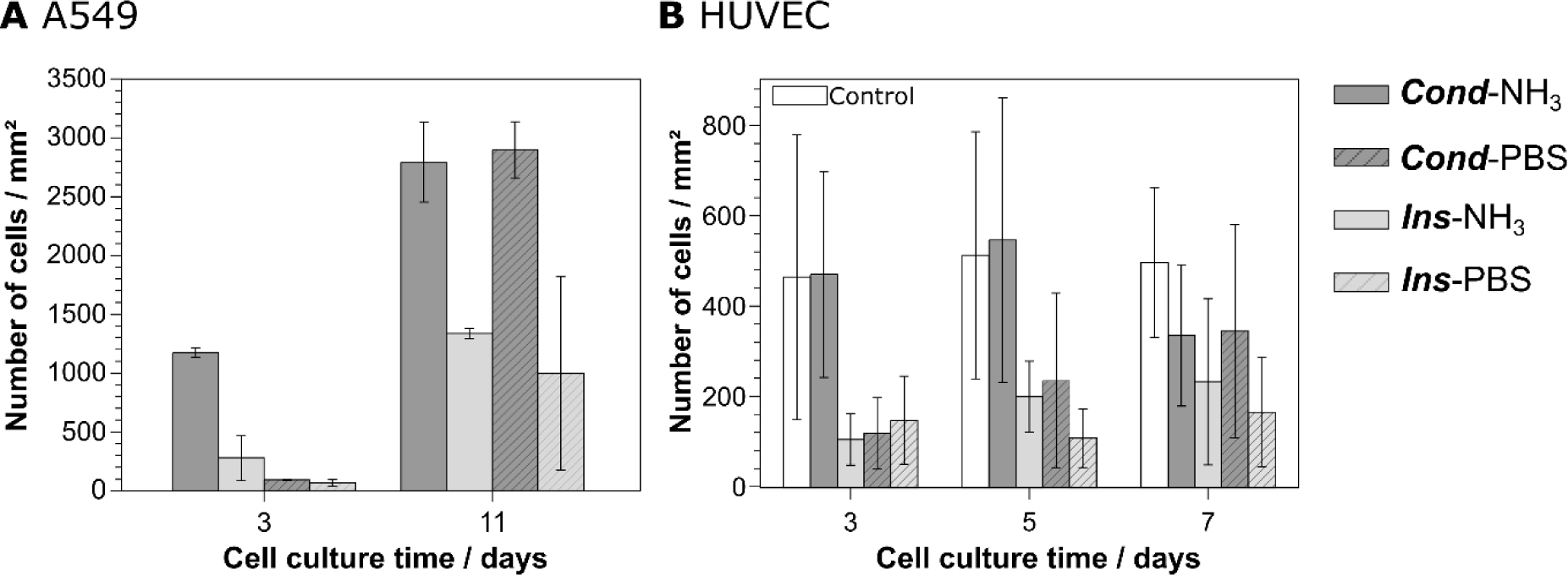
Ability of the materials to promote epithelium formation. Two epithelial cell lines (A549 cells and HUVECs) were seeded on the luminal surface. (**A**) A549 cell density at days 3 and 11 on each material’s lumen. Error bars: ± SD, (**B**) HUVECs density at day 3, 5 and 7 on each material’s lumen. Error bars: ± SD.

### Migration of hMSCs into the tubular scaffolds from the external surface

Materials were seeded with human mesenchymal stem cells (hMSCs) on their external surface and cultured for three weeks in order to evaluate their ability to be colonized. Cryosections stained with hematoxylin and eosin dyes (Figure 7) indicate that hMSCs deeply migrate in ***Cond*-NH_3_** at day 21, while only partial colonization is observed in other materials. Since porosity is poorly preserved in **-PBS**, cells lie on the top surface and do not migrate into the materials. Moreover, ***Ins*** materials seem less favorable for colonization compared to ***Cond*** ones, may be due to the looser collagen density within the walls observed at the TEM.

## 3. Conclusions

We describe here a strategy to replicate the morphology, the mechanics and the functions of native tubular tissues’ ECM using type I collagen exclusively.

Ice-templating has proven versatile as a shaping approach to fine-tune the materials porosity by adjusting the thermal boundary conditions imposed by the molds. This simple method allows orienting pores in either longitudinal or radial directions, providing a mean to tailor cell access to the scaffold. The topotactic fibrillogenesis conditions provide further control over the porosity while triggering the self-assembly of collagen molecules into biomimetic fibrils. Our materials offer cellular micro-environments closely resembling those of biological tissues, a lacking feature in materials currently used in the clinic.

Using exclusively type I collagen and in the absence of any cross-linkers, the elastic properties, at both the tissue and cellular length scales, reach those of native tissues. Control over the porosity and orientation given by our innovative approach directs the anisotropic *versus* isotropic character of the materials’ elastic properties, offering the opportunity to mimic mechanical behaviors of a wide range biological tissues.

Interactions between the materials and different cell lines − ECs and MSCs − demonstrate their potential to serve as biologically relevant substrates. On the basis of hMSCs migration experiments, ammonia fibrillogenesis proves more suitable to preserve pores and to facilitate cell colonization, ensuring a successful graft implantation. Epithelial and endothelial cells adhere well to the materials and proliferate until confluence is reached. ***Cond*** substrates appear to better support endothelialization at early time points compared to ***Ins*** surfaces, although this difference diminishes over time.

Overall, we introduce a new family of tubular collagen materials, exhibiting a unique combination of high concentration and porosity, with unprecedented resemblance to native tissues. Through their textural and compositional features, they promote both cell adhesion on the luminal surface and cell migration from the external surface, associated with mechanical performances suitable for functional replacement. We anticipate that these could prove central to a new range of grafts whose biomimetic features secure assimilation by the body of the patient.

Beyond their role as tissue grafts, our materials are expected to act as organotypic models of tubular tissues. These may answer a vast range of open questions — from cell-cell and cell-matrix interactions up to drug treatment response — bridging current gaps for improved patient care in respiratory, vascular, gastrointestinal and urinary tracts pathologies.

## 4. Experimental section

### 4.1. Fabrication of the collagen scaffolds

Collagen I was extracted from young rat tail tendons. After thorough cleaning of tendons with phosphate buffered saline (PBS; 1X) and 4 M NaCl, tendons were dissolved in 3 mM HCl. Differential precipitation with 300 mM NaCl and 600 mM NaCl, followed by redissolution and dialysis in 3 mM HCl, provided collagen of high purity. The final collagen concentration was determined using hydroxyproline titration with a spectrophotometer. To attain a high fibrillar collagen density in the model, the remaining collagen solutions were concentrated at 40 mg.mL^-1^. The solutions were then transferred into Vivaspin tubes with 300 kDa filter and centrifugated at 3000 g at 10°C, to reach this final concentration.

Concentrated collagen is introduced in between two cylindrical molds (inner mold: 3×5 mm; outer mold: 8×10 mm) whose ends are hermetically sealed by caps. For removal of air bubbles, sample is centrifuged at 1 rpm and10°C for 10 minutes. The sample is placed onto a home-made set-up that allows a continuous and regulated dipping of the sample in liquid nitrogen. It is composed of: 2 optical axes, 4 linear rod rail support guides, a lead screw, a DC motor (12 V), a speed controller, a transformer and an ABS 3D-printed part linking the support guide and the lead screw (Figure S1).

The sample is ice-templated at a dipping speed of 10 mm.min^-1^. Once frozen, the outer mold is quickly removed and a topotactic fibrillogenesis condition is applied. At low temperature, two conditions are compared: (a) exposure to ammonia vapors or (b) PBS (10X: 1.37 M, KCl 26.8 mM, Na_2_HPO_4_ 80.7 mM, and NaH_2_PO_4_ 14.7 mM). Pre-fibrillogenesis for (a) consists of exposing the sample to ammonia vapors at 0°C in an ice-water bath for 48 h, followed by removal of ammonia using distilled water vapors for 24 h at 37°C in a heat room. For (b), the sample is simply maintained in PBS (10X) at -3°C in a thermostatic bath for 72 h. For both (a) and (b), fibrillogenesis is completed by keeping the sample in PBS (5X: NaCl 685 mM, KCl 13.4 mM, Na_2_HPO_4_ 40.35 mM, and NaH_2_PO_4_ 7.35 mM) for 14 days at room temperature.

### 4.2. Scanning electron microscopy

After ice-templating, the frozen matrices were freeze-dried for 24h, and the resulting dried foams were cut into different sections to determine crystallization mechanism and pores alignement of the scaffold. Two sections were cut horizontally (perpendicular to the ice front growth direction), at the top and bottom. Two sections were cut longitudinally (parallel to the ice front growth direction), to depict the lumen and the intima-adventitia-like layers of the tubular construct. Samples were imaged using scanning electron microscopy (SEM), with prior sputter coating with a 10nm layer of gold. Observations were performed at 10 kV at various levels of magnification using a Hitachi S-3400N microscope.

### 4.3. Porosity quantification

Porosity was quantified using ImageJ Fiji (Figure S2). Images were first binarized by adjusting the threshold for transversal sections or using the *LabKit segmentation* plugin for the surfaces. Porosity was quantified using the ImageJ plugin *Analyze particles* which detects particles or round structures in images by analyzing intensity variations and identifying groups of pixels with high gradient values. Feret diameter (MaxFeret), minimum Feret diameter (MinFeret), aspect ratio and area were the set measurements. The Feret diameter corresponds to the diameter of the smallest circle enclosing the pore, whereas MinFeret corresponds to the diameter of the biggest circle enclosed within the pore. Data are expressed as mean ± SE (standard error). Pore orientation was quantified using the ImageJ plugin *OrientationJ*. Pore surface coverage was determined as the percentage of accessible pores on a given surface.

### 4.4. Confocal microscopy

After fibrillogenesis, semi-thin sections (200 µm-thick) were cut. Collagen was labelled with fluorescein isothiocyanate (FITC, 0.05 mg/mg of collagen) at 4°C overnight. Cellularized samples with A549 cells were fixed in between a glass slide and a coverslip with a handmade set up to maintain it, and imaged at the end of the culture. Observations were conducted on using a Zeiss LSM 980 upright confocal microscope.

### 4.5. Transmission electron microscopy

Hydrated samples were cross-linked with 2.5% paraformaldehyde (PFA), 2% glutaraldehyde, 0.18M sucrose and 0.1% picric acid for 12 h. The samples were subsequently post-fixed with uranyl acetate in ethanol for 12h, and dehydrated using baths with increasing concentrations of ethanol. Samples were incorporated in SPURR-S resins prior to sectioning. 70 nm ultrathin sections (Leica microtome) were contrasted with uranyl acetate and observed on a transmission electron microscope (FEI Tecnai Spirit G2) operating at 120 kV to observe the ultrastructural collagen features. Images were recorded on a Gatan CCD camera.

### 4.6. Polarized light microscopy

Samples were incorporated in 2,5% agarose and 200 µm semi-thin sections were cut with a vibratome (Compresstome), and observed under a Nikon Eclipse E600Pol microscope equipped with 4X, 10X and 40X objectives, a DXM 1200 CCD camera, and cross-polarizers to observe the birefringence of the sections.

### 4.7. Axial and circumferential Young’s modulus measurement

Uniaxial traction testing of porous collagen tubes was performed on a homemade stage with a 10 N load cell (ME-Meßsysteme GmbH, Germany). The tubes were first cut open and unfolded toward a flat geometry. Rectangular strips measuring 2 cm × 1 cm were then cut from the samples in both circumferential and longitudinal directions. To measure the thickness of the samples, a compression rheometer (MCR 302. Anton Paar) was used: the base of a cylindrical probe was approached at a constant speed toward the sample until a force was detected, corresponding to the thickness of the sample. For the uniaxial tests, each strip was held in place with serrated jaws blocking the ends. The initial distance between the jaws was adjusted until each strip was flattened with non-zero force measured, and this distance was set as the unloaded sample length. To ensure consistency and accuracy of the results, two pre-conditioning cycles were performed to produce a consistent force-elongation curve for each strip. Each strip was subjected to cyclic testing, and data from the third cycle’s loading phase were used for analysis. The deformation rate, *k* = ^*v*^*L*_*L*_, with *v* being the deformation speed and *L* the unloaded sample length, was maintained for each samples at a constant value of 0.8%.This protocol was followed for all tests. The resulting stress/strain curves were used to calculate the Young’s modulus, and only the linear portion of these curves was utilized (Figure S5). All measurements were carried out under immersion conditions in PBS (1X) at the physiological temperature of 37°C. Three tubes for each condition were used for statistical relevance.

### 4.8. Compression elastic Young modulus

The compression elastic modulus (Young’s modulus) was assessed by AFM using a Nanowizard 4 BioAFM (PK/Bruker) mounted on a Nikon Eclipse Ti2-U inverted microscope equipped with PetriDish Heater. Due to the difference in stiffness between the PBS (10X) and NH_3_ fibrillated samples, two appropriately matched probes with different spring constants were used for the measurements (qp-BioAC-CI and PPP-FMAuD from NANOSENSORS^TM^). Cantilevers were calibrated using the thermal calibration method in PBS (1X) at 37°C. In order to locate the walls of our sample (i.e. to ensure we were not scanning within a pore), initial blind 2D scans were performed. Subsequently, appropriately sized scans were conducted in these identified regions (Figure S5). The Hertz model was applied to fit each force-indentation curve, and stiffness for each scan point was extracted using JPK data processing software. All measurements were made under immersion conditions in PBS (1X) at a physiological temperature of 37°C.

### 4.9. Tubular matrix cellularization

#### HUVECs

Human umbilical vein endothelial cells (HUVECs, Lonza) in passages 5-8 were cultured at 37°C and 5% CO_2_ in EGM2 cell medium (Lonza). Cells were detached from the adhering surface using TrypLE (Life Technologies, Carlsbad, USA). HUVECs were seeded on the luminal surface of 6 mm diameter discs of the collagen scaffolds (or PDMS coated with Collagen IV as control) at a density of 5,000 cells/sample and cultured for 3, 5 and 7 days. At the end of the culture, samples were fixed with 4% paraformaldehyde, permeabilized with 0.1% Triton X-100, and then incubated with DAPI (Sigma Aldrich) and Phalloidin 495 (LifeTechnologies) to visualize nuclei and the F-actin cytoskeleton. A series of images z-stacks were acquired at 1 µm intervals using an inverted microscope (Nikon Eclipse Ti-U, Japan) equipped with a CCD camera (Orca AG; Hamamatsu, Japan) and a motorized x-y stage equipped with a 20X Plan Apo λ Ph2 DM objective. Cell proliferation was quantified by counting the number of nuclei (based on the DAPI signal) using the FIJI plugin Cell counter.

#### Epithelialization with A549

Discs of 6 mm diameter were seeded on the luminal surface with 1’500 A549 cells (equivalent to 5300 cells/cm^2^), and then grown in DMEM containing 25 mM D-glucose, 10 mM HEPES, 23.8 mM NaHCO3, 1 mM sodium pyruvate, 2 mM L-glutamine, 100 U/ml penicillin, 100 µg/ml streptomycin, 10 µg/ml gentamycin, and 10% FCS in a 3% O_2_, 5% CO_2_ and 92% air atmosphere. The medium was changed every 48h hours.

Samples were fixed at days 3 and 11 after seeding with 4% formaldehyde in PBS (1X) for 16 hours at 4°C, then rinsed in PBS. After permeabilization with 0.1% Triton-X100 in PBS (1X) for 10 minutes and BSA-blockage in 3% BSA/0.1% Tween-20 in PBS, samples were stained with 1 µg/mL TRITC-conjugated phalloidin (Sigma) for 15 min at room temperature. They were rinsed several times in PBS (1X), stained with DAPI and placed in aqueous mounting medium (Citifluor AF1) for confocal microscope analysis.

#### External surface seeding with human mesenchymal stem cells (hMSC)

Six mm diameter punches were seeded on their external surface with 50’000 hMSCs (equivalent to 176’000 cells/cm2), and then grown for 3 weeks in DMEM containing 25 mM D-glucose, 10 mM HEPES, 23.8 mM NaHCO3, 1 mM sodium pyruvate, 2 mM L-glutamine, 100 U/ml penicillin, 100 µg/ml streptomycin, 10 µg/ml gentamycin, and 10% FCS in a 5% CO2 – 95% air atmosphere. The medium was changed every 48h hours.

After 3 weeks of culture, samples were fixed with 4% formaldehyde in PBS (1X) for 16 hours at 4°C, then rinsed in PBS. Samples were successively transferred in 15%, 30% sucrose in PBS (1X) and then in 50% OCT / 30% sucrose mixture at 4°C until equilibration. Finally, samples were progressively embedded in 100% OCT on dry ice and conserved at -80°C until cryosections were performed. Ultrathin cryo-sections (10 µm-thick) were cut and post-fixed in ice-cold acetone for 15 minutes. Samples were then stained with haematoxylin and eosin dies for 5 minutes and placed in aqueous mounting medium for microscope analysis.

## Supporting information

Supplementary information is available here.

## Acknowledgements

The authors kindly acknowledge the help from F. Lam and C. Chaumeton with confocal fluorescence microscopy. C. Djédiat is equally acknowledged for the preparation of samples for TEM observations. The authors gratefully acknowledge the contribution of C. Parisi and E. Budimirovic for their initial contributions on the subject.

This work was supported by Agence Nationale de la Recherche (ANR) under grant agreement ANR-20-CE19-0029. IM acknowledges the PhD fellowship from the Physics and Chemistry of Materials’ graduate school (ED397).

## Notes

### Competing Interest Statement

The authors have declared no competing interest.

## References

[1] M. Carrabba, P. Madeddu, Front Bioeng Biotechnol 2018, 6, 1.

[2] L. Cen, W. Liu, L. Cui, W. Zhang, Y. Cao, Pediatr Res 2008, 63, 492.

[3] M. M. Giraud Guille, N. Nassif, F. M. Fernandes, In Materials design inspired by nature: function through inner architecture (Eds.: Fratzl, P.; W.C. Dunlop, J.; Weinkamer, R.), The Royal Society of Chemistry, 2013, pp. 107–127.

[4] C. B. Weinberg, E. Bell, Science (1979) 1986, 231, 397.

[5] J. Hirai, T. Matsuda, Cell Transplant 1996, 5, 93.

[6] X. Li, J. Xu, C. T. Nicolescu, J. T. Marinelli, J. Tien, Tissue Eng Part A 2017, 23, 335.

[7] C. E. Ghezzi, B. Marelli, N. Muja, S. N. Nazhat, Acta Biomater 2012, 8, 1813.

[8] V. A. Kumar, J. M. Caves, C. A. Haller, E. Dai, L. Liu, S. Grainger, E. L. Chaikof, Acta Biomater 2013, 9, 8067.

[9] C. Parisi, B. Thiébot, G. Mosser, L. Trichet, P. Manivet, F. M. Fernandes, Biomater Sci 2022, 10, 6939.

[10] M. Madaghiele, A. Sannino, I. V. Yannas, M. Spector, J Biomed Mater Res A 2008, 85, 757.

[11] K. M. Pawelec, A. Husmann, R. J. Wardale, S. M. Best, R. E. Cameron, J Mater Sci Mater Med 2015, 26, 91.

[12] M. J. W. Koens, A. G. Krasznai, A. E. J. Hanssen, T. Hendriks, R. Praster, W. F. Daamen, J. A. van der Vliet, T. H. van Kuppevelt, Organogenesis 2015, 11, 105.

[13] J. F. Cavallaro, P. D. Kemp, K. H. Kraus, Biotechnol Bioeng 1994, 43, 781.

[14] L. Magnan, G. Labrunie, M. Fénelon, N. Dusserre, M. P. Foulc, M. Lafourcade, I. Svahn, E. Gontier, J. H. Vélez V., T. N. McAllister, N. L’Heureux, Acta Biomater 2020, 105, 111.

[15] S. Deville, Adv Eng Mater 2008, 10, 155.

[16] K. Qin, C. Parisi, F. M. Fernandes, J Mater Chem B 2021, 9, 889.

[17] M. C. Gutiérrez, M. L. Ferrer, F. Del Monte, Chemistry of Materials 2008, 20, 634.

[18] K. M. Pawelec, A. Husmann, S. M. Best, R. E. Cameron, Appl Phys Rev 2014, 1, 021301.

[19] C. Rieu, C. Parisi, G. Mosser, B. Haye, T. Coradin, F. M. Fernandes, L. Trichet, ACS Appl Mater Interfaces 2019, 11, 14672.

[20] E. Thibert, F. Dominé, Journal of Physicial Chemistry B 1997, 101, 3554.

[21] J. Schindelin, I. Arganda-Carreras, E. Frise, V. Kaynig, M. Longair, T. Pietzsch, S. Preibisch, C. Rueden, S. Saalfeld, B. Schmid, J. Y. Tinevez, D. J. White, V. Hartenstein, K. Eliceiri, P. Tomancak, A. Cardona, Nat Methods 2012, 9, 676.

[22] D. A. Knight, S. T. Holgate, Respirology 2003, 8, 432.

[23] Y. Bouligand, Comptes Rendus Chimie 2008, 11, 281.

[24] F. Gobeaux, G. Mosser, A. Anglo, P. Panine, P. Davidson, M. M. Giraud-Guille, E. Belamie, J Mol Biol 2008, 376, 1509.

[25] M. M. Giraud-Guille, E. Belamie, G. Mosser, C. Helary, F. Gobeaux, S. Vigier, Comptes Rendus Chimie 2008, 11, 245.

[26] M. F. Ashby, Y. J. M. Bréchet, Acta Mater 2003, 51, 5801.

[27] P. J. Critser, S. T. Kreger, S. L. Voytik-Harbin, M. C. Yoder, Microvasc Res 2010, 80, 23.

