## Supplementary information is available here. for "Tunable biomimetic materials elaborated by ice templating and self-assembly of collagen for tubular tissue engineering"

*by*

*Isabelle Martinier[1], Florian Fage[1], Alshaba Kakar[1], Alessia Castagnino[2], Emeline Saindoy[3], Joni Frederick[2], Ilaria Onorati[3], Valérie Besnard[3], Abdul I. Barakat[2], Nicolas Dard[3], Emmanuel Martinod[3], Carole Planes[3], Léa Trichet[1]*, Francisco M. Fernandes[1]**

[1] Laboratoire de Chimie de la Matière Condensée de Paris, Sorbonne Université, 4 place Jussieu, 75005 Paris, France

[2] LadHyX, CNRS, Ecole Polytechnique, Institut Polytechnique de Paris, Palaiseau, France

[3] Laboratoire Hypoxie & Poumon, Assistance Publique–Hôpitaux de Paris, Hôpitaux Universitaires Paris Seine-Saint-Denis, Hôpital Avicenne, Chirurgie Thoracique et Vasculaire, Université Paris 13, Sorbonne Paris Cité, UFR Santé, Médecine et Biologie Humaine, Bobigny, France

**Table S1**. The different pairs of mold materials and their corresponding thermal conductivity differences used for the study of the interface position during ice-templating of a collagen solution. The difference was calculated by subtracting the inner material thermal conductivity from the outer one. Interface position is given in percentage as the distance to the lumen relative to the total wall thickness. The materials include aluminum (237 W/(m.K)), brass (110 W/(m.K)), stainless steel (45 W/(m.K)), acrylonitrile butadiene styrene (ABS) (0.17 W/(m.K)) and high impact polystyrene (HIPS) (0.11 W/(m.K)).

| Number | Outer material | Inner material | Thermal conductivity difference (W/(m.K)) | Interface position (%) |
|---|---|---|---|---|
| 1 | Stainless steel | Aluminum | -192 | 62.5 |
| 2 | Brass | Aluminum | -127 | 38.1 |
| 3 | Brass | Brass | 0 | 27.9 |
| 3 | Stainless steel | Stainless steel | 0 | 17.6 |

| | | | | |
|---|---|---|---|---|
| 4 | Stainless steel | ABS | 44.8 | 16.4 |
| 4 | Stainless steel | HIPS | 44.9 | 18.8 |
| 5 | Brass | Stainless steel | 65 | 13.9 |
| 6 | Brass | ABS | 109.8 | 8.5 |
| 6 | Brass | HIPS | 109.9 | 5.6 |

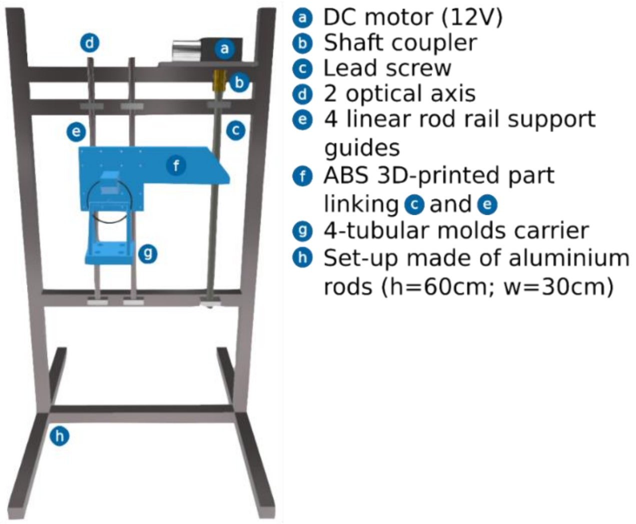


**Figure S1.** Homemade set-up for unidirectional controlled ice-templating.

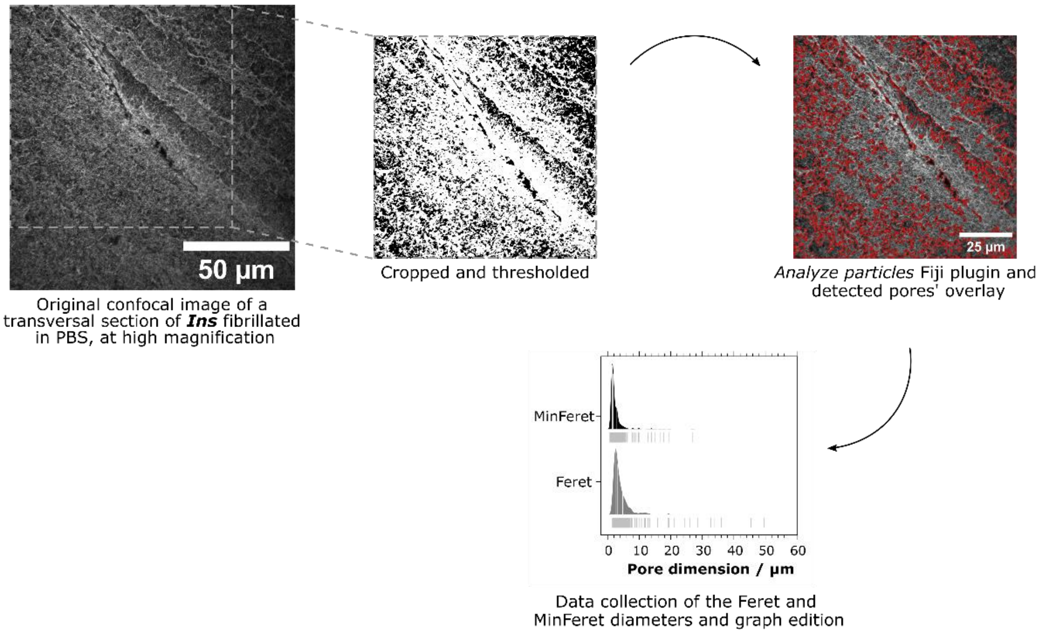


**Figure S2.** Pores dimensions analysis using ImageJ Fiji of ***Ins*** scaffold fibrillated in PBS (10X) and imaged by confocal microscopy at high magnification.

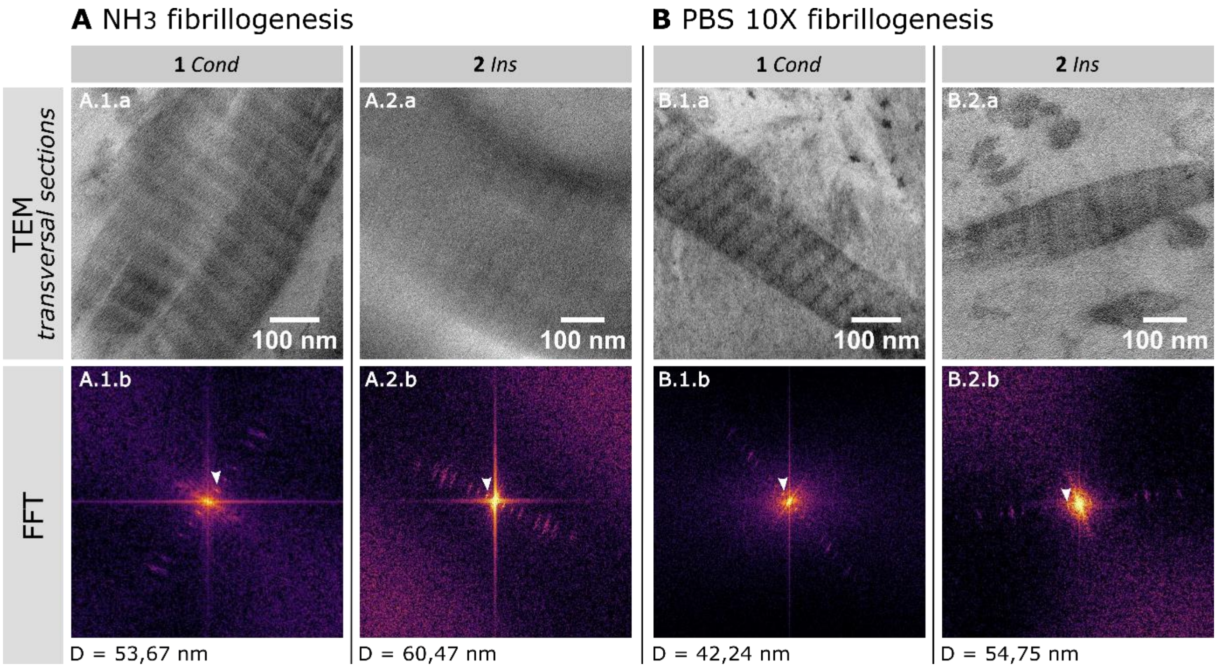


**Figure S3.** (A.1.a, A.2.a, B.1.a, B.2.a): High magnification of the fibrils' striation for each scaffold, imaged by transmission electron microscopy. (A.1.b, A.2.b, B.1.b, B.2.b): Corresponding fast Fourier transforms (FFT) and the measured D-bandings.

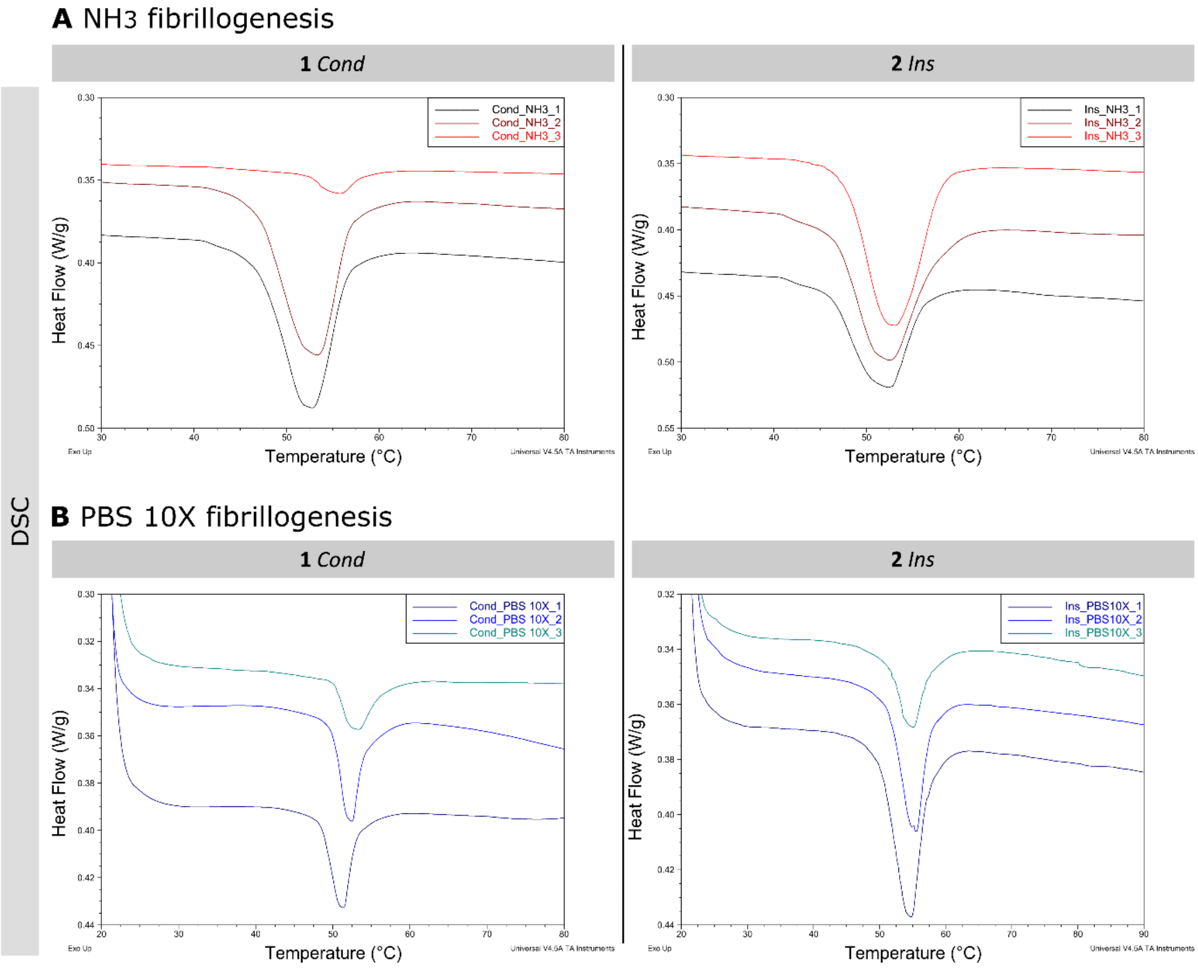


**Figure S4.** Detailed thermograms of the denaturation temperature measurements for each scaffold. Samples were heated from 20 to 120°C at a rate of 5°C.min$^{-1}$)

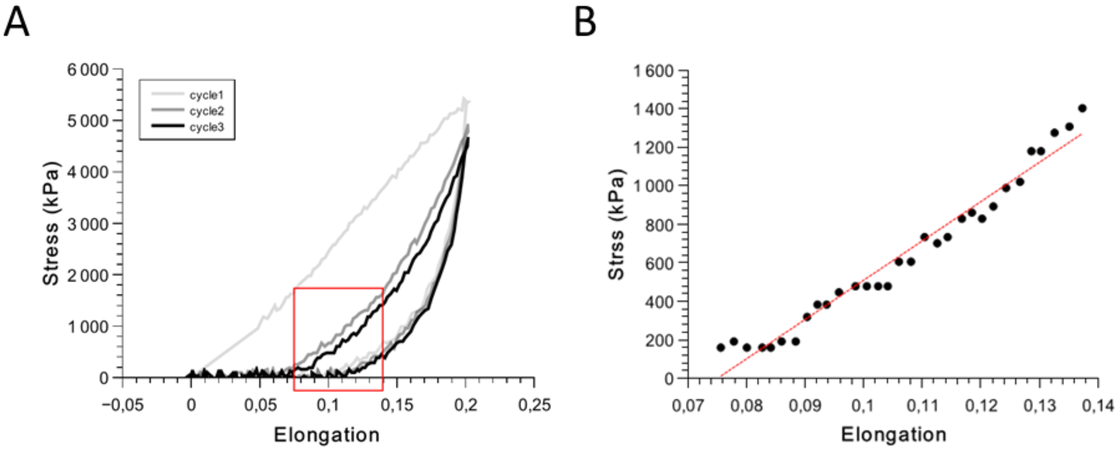


**Figure S5.** (**A**) Representative stress/strain curves for traction cycles. Each cycle is color-coded with Cycle 1 shown in light grey, Cycle 2 in grey, and Cycle 3 in black. (**B**) Illustration of stress-strain fitting for the linear region of the third cycle.

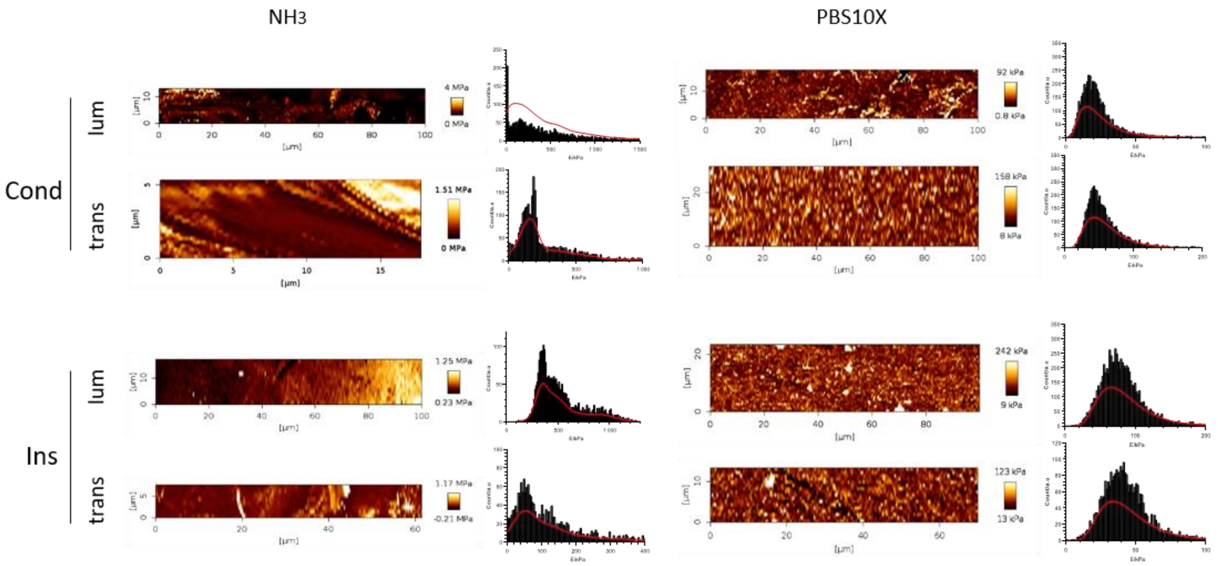


**Figure S6.** Typical Young's modulus maps extracted from AFM measurements and their corresponding statistical distributions. The distributions are smoothed using the kernel density estimation method (in red).
